# Physical geography, isolation by distance and environmental variables shape genomic variation of wild barley (*Hordeum vulgare* L. ssp. *spontaneum*) in the Southern Levant

**DOI:** 10.1101/2021.09.15.460445

**Authors:** Che-Wei Chang, Eyal Fridman, Martin Mascher, Axel Himmelbach, Karl Schmid

## Abstract

Determining the extent of genetic variation that reflects local adaptation in crop wild relatives is of interest to discovering useful genetic diversity for plant breeding. We investigated the association of genomic variation with geographical and environmental factors in wild barley (*Hordeum vulgare L. ssp. spontaneum*) populations of the Southern Levant using genotyping-by-sequencing (GBS) of 244 accessions of the Barley1K+ collection. Inference of population structure resulted in four genetic clusters that corresponded to eco-geographical habitats and a significant association of lower gene flow rates with geographical barriers, e.g. the Judaean Mountains and the Sea of Galilee. Redundancy analysis (RDA) revealed that spatial autocorrelation explained 45% and environmental variables explained 15% of total genomic variation. Only 4.5% of genomic variation was exclusively attributed to environmental variation if the component confounded with spatial autocorrelation was excluded. A synthetic environmental variable combining latitude, solar radiation, and accumulated precipitation explained the highest proportion of genomic variation (3.9%). After correcting for population structure, soil water capacity was the most important environmental variable explaining 1.18% of genomic variation. Genome scans with outlier analysis and genome-environment association studies were conducted to identify signatures of adaptation. RDA and outlier methods jointly detected selection signatures in the pericentromeric regions of chromosome 3H, 4H, and 5H, but they mostly disappeared after correction for population structure. In conclusion, adaptation to the highly diverse environments of the Southern Levant over short geographical ranges has a small effect on the genomic diversity of wild barley highlighting the importance of non-selective forces in genetic differentiation.

## Introduction

Local adaptation is an essential strategy for plants to survive in stressful environments because they are sessile. Natural selection in heterogeneous environments leads to higher fitness of local genotypes, but gene flow can offset genetic differentiation resulting from local adaptation and reduce fitness (Kawecki and Ebert 2004). In addition, genetic drift and demographic history contribute to genetic differentiation and confound adaptive variation with neutral variation (Günther and Coop 2013; Kawecki and Ebert 2004; López-Goldar and Agrawal 2021). Therefore, the combination of selective and nonselective forces simultaneously shapes genetic variation and leads to geographic patterns of population divergence and allele frequency distribution. Determining how different population genetic processes affect the geographic distribution of genetic variation is a key component in the study of plant adaptation. Investigating the role of adaptive and non-adaptive processes on genomic variation is of particular interest for wild relatives of crop plants, as this may allow the discovery of useful genetic variation for plant breeding (Turner-Hissong et al. 2020).

Wild barley (*Hordeum vulgare L. ssp. spontaneum*) is a very suitable model species to study local adaptation of crop wild relatives, as it occurs over a wide geographical range in the Fertile Crescent (Harlan and Zohary 1966) and Eastern and Central Asia (Dai et al. 2012). Within this range, wild barley in the Levant occupies very heterogeneous environments within a short geographical distance (Hübner et al. 2009; Nevo et al. 1979; Volis et al. 2001), and its genetic diversity is much higher than in other areas of the Fertile Crescent (Jakob et al. 2014; Pankin et al. 2018; Russell et al. 2016). Wild barley populations from the southern Levant show a strong correlation of genetic and environmental distances (Hübner et al. 2009). Population structure reflects eco-geographic habitats (Hübner et al. 2012, 2009) and distinguishes between northern and southern genetic clusters correlated with latitude and precipitation gradients (Jakob et al. 2014; Russell et al. 2016). With common garden experiments, previous studies revealed that eco-geography is correlated to morphological traits (Hübner et al. 2013), phenotypic plasticity (Galkin et al. 2018), and rhizosphere microbiota (Terrazas et al. 2020).Moreover, transplantation experiments showed a correlation between the geographic origin of wild barley ecotypes and fitness in different environments, suggesting local adaptation (Volis 2011; Volis et al. 2002a,b). In addition to a broad geographic scale, environmental differences at a fine geographic scale also contribute to genetic diversification in wild barley (Bedada et al. 2014; Nevo et al. 2005; Wang et al. 2018). Overall, these results suggest a strong relationship between environmental differences, genetic divergence and phenotypic diversity of wild barley populations, supporting the hypothesis of local adaptation of wild barley in the southern Levant. However, the relative contributions of environmental and non-selective forces to genetic variation and the genetic architecture of adaptive traits remain mostly unclear due to the lack of appropriate statistical approaches, fine-scale environmental data and sufficient genome-wide markers.

Wild barley serves as a valuable genetic resource for barley breeding because domestication and modern breeding have greatly reduced the genetic diversity of cultivated barley (*Hordeum vulgare L. ssp. vulgare*; Caldwell et al. 2006; Kilian et al. 2006). Since wild barley shows no reproductive barrier with cultivated barley (Nevo et al. 1979), the genetic diversity of cultivated barley can be expanded by introducing alleles from wild populations (Dawson et al. 2015). Numerous studies have shown evidence of local adaptation of wild barley to different environments (Bedada et al. 2014; Galkin et al. 2018; Hübner et al. 2013; Nevo et al. 1979; Volis 2011; Volis et al. 2002a,b, 2004; Wang et al. 2018), therefore wild barley is expected to have considerable genetic variation that contributes to adaptation to various abiotic stresses (Dawson et al. 2015). Wild barley has been repeatedly used as a source of novel alleles to improve stress tolerance in barley breeding (Baum et al. 2003; Pham et al. 2019), but widespread use of wild barley has been limited due to its large genome size and undesirable traits (Schmid et al. 2018).

Genotyping-by-sequencing (GBS; Elshire et al. 2011; Poland et al. 2012) and high-quality barley genome assembly (Jayakodi et al. 2020; Mascher et al. 2017), enable the exploration of genomic variation under environmental selection and the search for useful genetic variation in wild barley. In this study, we aimed to (1) describe the population structure of wild barley from the southern Levant and place it in the context of a worldwide sample (Milner et al. 2019), (2) examine geographic patterns of gene flow in the southern Levant, (3) characterize the relative contributions of environmental gradients and space to genomic variation and population structure, and (4) identify putative adaptive loci. Overall, our results indicate the greater importance of geography and spatial autocorrelation than selection for local adaptation in shaping genomic variation in wild barley in the southern Levant, although diverse environments, particularly water availability, show significant associations with genetic differentiation.

## Materials and Methods

### Plant material and genotyping by sequencing

We genotyped 244 wild barley accessions collected in the southern Levant region. These accessions, hereafter named as B1K+ accessions, include 191 accessions from Barley 1K (B1K) collection (Hübner et al. 2009) and 53 accessions from an unpublished collection named as HOH, collected in 2005, 2009, and 2011 by K.S. (Figure S1; File S1). The GBS library was constructed using genomic DNA digested with the restriction enzyme *PstI* and *MspI* and the processes as described by Milner et al. (2019). In addition, published GBS data of 1,121 wild barley accessions from IPK seed bank (Milner et al. 2019) was included as a part of our work. Because IPK genebank includes accessions from Israel, we specify the source of accessions by mentioning accessions from Israel that belong to IPK in the article to avoid confusion. Identification of single-nucleotide polymorphism (SNP) was performed as Milner et al. (2019). The detailed workflow of genotypic data filtration is described in Supplementary Information and summarized by Figure S2.

### Environmental data

To investigate the relationship between genetic variation and environmental gradients, we used environmental data including (1) climate data from the *WorldClim2* database (Fick and Hijmans 2017) with a resolution of 30 arcseconds [~1 km], (2) soil data from the *SoilGrids* database (Hengl et al. 2017) with a resolution of 250m, (3) topographic variables based on elevation data from the *SRTM* database (https://srtm.csi.cgiar.org/) with a resolution of 90m, and (4) geographic coordinates of collection points (Supplementary Information; File S1). To mitigate the problem of collinearity for redundancy analysis (RDA; Legendre and Legendre 2012), highly correlated environmental variables were grouped by hierarchical clustering with a customized clustering index (Supplementary Information; Figure S3), and the first principal components of the grouped variables were used as synthetic variables. Next, all environmental variables, including the synthetic variables, were selected based on variance inflation factors (VIFs) until all VIFs were less than 5. The details of this process are described in the Supplementary Information. Finally, 12 environmental variables, including 7 synthetic variables and 5 nonsynthetic variables (Table S1 and S2), were selected for environmental association analyses.

### Analysis of population structure

The number of ancestors and ancestry coefficients were estimated using the model-based method *ALStructure*. (Cabreros and Storey 2019). *ALStructure* uses a likelihood-free algorithm to derive estimates through minimal model assumptions and is generally superior to existing likelihood-based methods in terms of accuracy and computational speed (Cabreros and Storey 2019). The method does not assume Hardy-Weinberg equilibrium within populations, but defines the number of ancestral populations (*K*) as the rank of a matrix consisting of individual-specific allele frequencies (Leek 2011). Optimal *K* was calculated using the *estimate_d* function of the R package *alstructure* (Cabreros and Storey 2019), and ancestry coefficients were estimated using the *alstructure* function. A range of *K* values, from 2 to 8, was also used to examine stratification of population structure. In addition to *ALStructure*, principal component analysis (PCA) and neighbor-joining (NJ) were also performed. Missing genotypic values (~3% of the dataset) were replaced by the average number of alternative alleles at each SNP locus before performing PCA.

To analyze genetic differentiation, we calculated *F_ST_* and Nei genetic distance between genetic clusters defined by *ALStructure*. Accessions were assigned to genetic clusters according to the highest ancestry coefficient calculated by *ALStructure* with the optimal value of *K*. The *F_ST_* values were calculated as the ratio of the average values according to Bhatia et al. (2013) and the genetic distances of Nei were calculated using the function *stamppNeisD* of the R package *StAMPP* (Pembleton et al. 2013).

### Investigation of gene flow pattern

To discover gene flow barriers that may explain observed population structure, analysis of estimated effective migration surfaces (*EEMS;* Petkova et al. 2016) was conducted. To perform *EEMS*, B1K+ accessions were clustered into 58 demes corresponding to the locations of collection sites. *EEMS* was conducted in three independent runs of Markov chain Monte Carlo (MCMC), and the results of three runs were averaged. Each MCMC chain included 10 million burn-in iterations and 10 million post-burn-in iterations thinned by an interval of 5000 iterations. Outputs of *EEMS* were processed with the R package *rEEMSplots*. To examine whether geographical barriers contribute to genetic isolation, we classified map pixels into barriers and non-barrier pixels according to geographical elevation (details in Supplementary Information; Figure S4) and conducted a Wilcoxon test to inspect the hypothesis that geographical barriers are significantly associated with lower gene flow rates.

As a complementary method of *EEMS*, unbundled principal components (unPC; House and Hahn 2018) was employed to reveal potential long-distance migration. *unPC* scores, a ratio of PCA-based genetic distance on population-level to geographical distance between demes, were computed with the R package *unPC*. Original *unPC* scores were transformed by Box-Cox transformation to an approximate Gaussian distribution, and a Student’s t-test with two-tailed significance level of 0.05 was performed to identify extreme comparisons.

To infer asymmetric gene flows, we implemented a coalescent-based inference (CBI) method described by Lundgren and Ralph (2019). We manually grouped accessions into ten geographical regions (Figure S5) such that each region covered roughly equal geographical areas as suggested by Lundgren and Ralph (2019) and also considered the gene flow pattern inferred by *EEMS.* Sample sizes in each region ranged from 10 to 41 with an average of 24.4. Next, we created an adjacency matrix to allow gene flow between adjacent regions (Figure S5). As the input for CBI, pairwise genetic distances were computed as the average number of different alleles across SNPs. CBI was conducted by using the R package *gene.flow.inference* (Lundgren and Ralph 2019) with 2 million pre-burn-in iterations, 60 million burn-in iterations, and 100 million post-burn-in iterations followed by a thinning process for every 5,000 iterations to rule out serial correlations. Medians of gene flow rates and coalescence rates were computed from posterior distributions and 95% credible intervals were calculated with the highest density interval method by using the R package *bayestestR* (Makowski et al. 2019).

### Partitioning genomic variation

To partition genomic variation into components explained by different factors, we conducted redundancy analysis (RDA), a multivariate method for studying a linear relationship of two or several matrices (Legendre and Legendre 2012). Specifically, we used simple RDA and RDA conditioned on covariates, i.e., partial RDA, to estimate the proportion of SNP variation explained by environmental variables, spatial autocorrelation, and population structure. RDA was performed with the function *rda* of the R package *vegan* (Oksanen et al. 2019). For all RDA models in this study, we performed 5,000 permutations to test the significance of explanatory variables with the R function *anova.cca.*

To model the effect of spatial autocorrelation on SNP variation, distance-based Moran’s eigenvector maps (dbMEMs) were used in RDA (Dray et al. 2006; Legendre and Legendre 2012). First, a network of 58 collection sites was built with the Gabriel graph, and a spatial weighting matrix of inverse geographical distances (*km*^-1^) was constructed accordingly by following the method of Forester et al. (2018). Next, the spatial weighting matrix was decomposed to generate dbMEMs. Subsequently, forward selection was performed to identify dbMEMs that significantly associate with spatial genetic structure by using *forward.sel* function (Dray et al. 2019). The selected dbMEMs with positive and negative eigenvalues, corresponding to broad-scale and fine-scale spatial structures, were both used in RDA to capture comprehensive spatial autocorrelation. To represent population structure in RDA, ancestry coefficients estimated by *ALStructure* with the optimal *K* values were used. We fitted RDA models by replacing SNP data with ancestry coefficients as response variables to quantify relative contributions of environments and spatial autocorrelation to population structure.

To evaluate the effects of individual environmental variables on SNP variation, we sequentially fitted one environmental variable as an explanatory variable at a time and treated ancestry coefficients as covariates in RDA models. Considering the correlation between environmental variables (Figure S3 C), we conducted additional permutation tests for marginal effects of environmental variables in a model including all the environmental variables by setting the parameter *by* = ‘*margin*’ for *anova.cca.* This method tests the significance of each environmental variable while excluding the effect that confounds with the other environmental variables.

### Linkage disequilibrium

Linkage disequilibrium (LD) was evaluated as pairwise *r*^2^ of SNPs by using *snpgdsLDMat* of the R package *SNPRelate* (Zheng et al. 2012) with a window size of 250 markers. To evaluate genome coverage of markers, genome-wide LD decline against physical distance was fitted by using local polynomial regression and the formula of Hill and Weir (1988). Local polynomial regression was carried out by using the R function *loess* with a smoothing parameter of 0.005.

### Identification of selection signatures

As a genome-environment association (GEA) method, RDA has high detection power and a low false-positive rate in identifying adaptation signatures (Capblancq et al. 2018; Forester et al. 2016, 2018). Thus, we scanned SNPs by performing simple RDA and partial RDA. Simple RDA was done by treating 27,147 SNPs as response variables and twelve environmental variables as explanatory variables. To control false positives due to population structure, partial RDA was performed by using ancestry coefficients estimated with the optimal *K* as covariates. A statistical framework proposed by Capblancq et al. (2018) was used for statistical tests and controlling false discovery rates (FDR). Briefly, loadings of SNPs in the first four RDA axes, selected according to the proportion of explained variation (Figure S6), were converted into Mahalanobis distances that approximated to a chi-squared distribution with four degrees of freedom. Next, p-values and q-values were computed accordingly, and SNPs with FDR < 0.05 were considered as candidate adaptive SNPs. The statistical test was conducted with the R function *rdadapt* (Capblancq et al. 2018).

Besides RDA, the latent factor mixed model (LFMM; Caye et al. 2019), which an univariate GEA method, was performed by using the R package *lfmm* (Caye et al. 2019) with the parameter *K* = *4* to correct population structure, and q-values were subsequently computed. SNPs with FDR < 0.05 were considered as candidate adaptive SNPs.

As a complement to GEA methods, outlier SNPs with an extreme divergence between genetic clusters were detected by *X^T^X* statistics (Günther and Coop 2013). We assigned accessions to genetic clusters according to the highest ancestry coefficient estimated by *ALStructure* with the optimal *K* and calculated *X^T^X* by using *BAYPASS* ver2.1 (Gautier 2015). *BAYPASS* was run by setting 25 short pilot runs, 100,000 burn-in iterations and 100,000 post-burn-in iterations with a thinning interval of 40 iterations. A significance threshold of *X^T^X* was determined by the 99.5% quantile of pseudo-observed *X^T^X* (Gautier 2015) calculated from neutral markers simulated by *simulate.baypass* (Gautier 2015).

### Gene ontology enrichment

To investigate biological functions related to putatively adaptive loci, we conducted gene ontology (GO) enrichment analysis with gene annotations of the barley ‘Morex v2’ genome (Mascher 2019). Over-representation of GO terms for genes within 500 bp adjacent intervals of candidate SNPs was tested by Fisher’s exact test with 10,000 runs by using *SNP2GO* (Szkiba et al. 2014), and GO terms with FDR < 0.05 were regarded as significantly enriched. Annotations of genes within 500 bp upstream and downstream of the candidate SNPs were also reported.

## Results

### Summary of genotypic data

SNP calling and preliminary filtration resulted in 101,711 SNPs for 1,365 accessions, including 1,121 IPK accessions and 244 B1K+ accessions (Supplementary Information; Figure S2). Depending on the requirements of analyses, we selected different subsets from 101,711 SNPs as follows. For the joint population structure analysis of IPK and B1K+ accessions, we selected 4,793 geographically diverse SNPs that are polymorphic (minor allele frequency; MAF ≥ 0.05) among 72 IPK accessions originating in 13 countries (Russell et al. 2016). This joint dataset is LD-pruned and has a missing proportion of 0.043 (Supplementary Information; Figure S2). For analyses of B1K+ accessions, we selected 58,616 SNPs with an overall missing proportion of 0.029 and maximal individual missing proportion of 0.059, and further filtration resulted in 19,601 SNPs (LD-pruned; MAF ≥ 0.01) and 27,147 SNPs (unpruned; MAF ≥ 0.05; Details in Supplementary Information; Figure S2)

### Population structure and spatial genetic pattern

Inference of population structure among B1K+ accessions with *ALStructure* (Cabreros and Storey 2019) identified four clusters (Figure 1 A and B), corresponding to the Mediterranean northern region, semi-arid coastal region, Judaean Desert, and Negev Desert (Figure 1C). Hereafter, we named the four B1K+ clusters as North, Coast, Eastern Desert, and Southern Desert. With *K = 4*, 174 of 244 (71.3 %) accessions have a highest ancestry coefficient less than 0.9. The first three principal components (PCs) represented the clusters corresponding to the *ALStructure* results (Figure 1 A and B). On the first PC axis, the northern cluster separated from two desert clusters, and on the second PC axis, the coastal cluster separated from the others. On the third PC axis, the southern desert cluster separated from the eastern desert cluster. The three PC axes explained 4.73%, 3%, and 2.83% of the variation, respectively. A hierarchical population structure was evident in the NJ tree (Figure 2A) and in the *ALStructure* analysis with K = 2-8 (Figure 2B). To evaluate the importance of marker density and additional samples to population structure analysis, in addition to original dataset, we performed PCA and *ALStructure* by either including or removing the HOH accessions with random selection of 100 and 5,000 SNPs. The dataset with 100 SNPs did not allow to identify genetic clusters while datasets with 5,000 SNPs separated into four genetic clusters by using the first four PCs even with the sample of original B1K accessions without the HOH accessions. However, *ALStructure* could only identify three ancestral populations (*K* = 3) if the HOH accessions were excluded (File S2).

**Figure 1.**
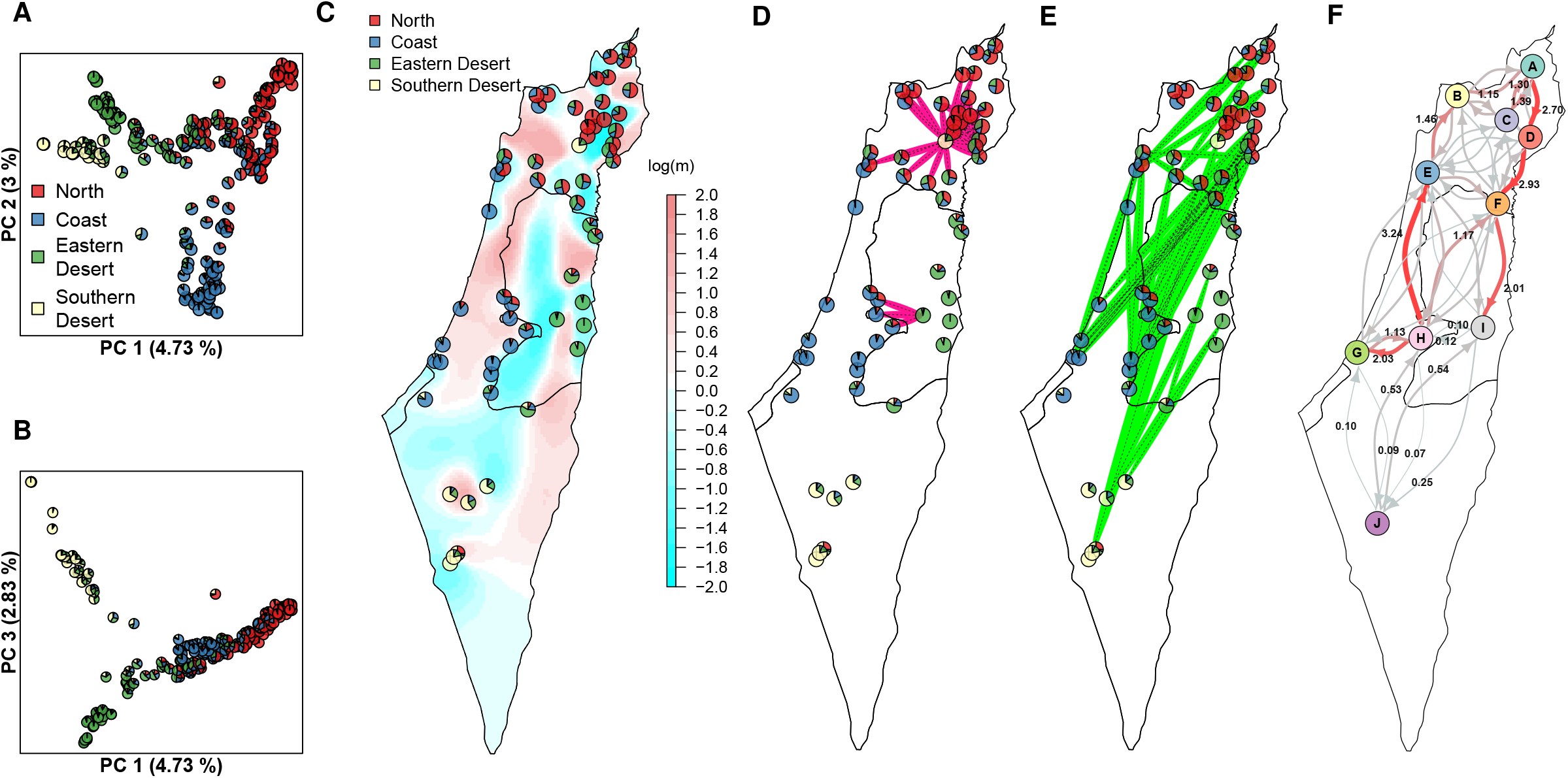
Spatial genetic structure of 244 B1K+ accessions and results of gene flow analysis. (A) PCA plot of the first and second PC axes. (B) PCA plot of the first and third PC axes. Pie charts in PCA plots represent ancestry coefficients of individuals estimated by *ALStructure* with *K* = *4*. (C) Distribution of genetic clusters and effective migration surface. Average ancestry coefficients of individuals in collection sites are shown by pie charts. Color gradient represents gene flow rates estimated by *EEMS*. (D) Comparisons with *unPC* scores higher than the top 2.5% threshold that indicates a significantly low genetic similarity over a short geographical distance. (E) Comparisons with *unPC* scores lower than the bottom 2.5% threshold that indicates a significantly high genetic similarity over a long geographical distance. (F) Gene flow rates inferred by the coalescent-based inference method, representing the probabilities per unit of time that individuals in a region *i* are descended from a region *j* (Lundgren and Ralph 2019). The thickness of arrows and the depth of red color is proportional to gene flow rates. The complete result is shown in Table S3.

**Figure 2.**
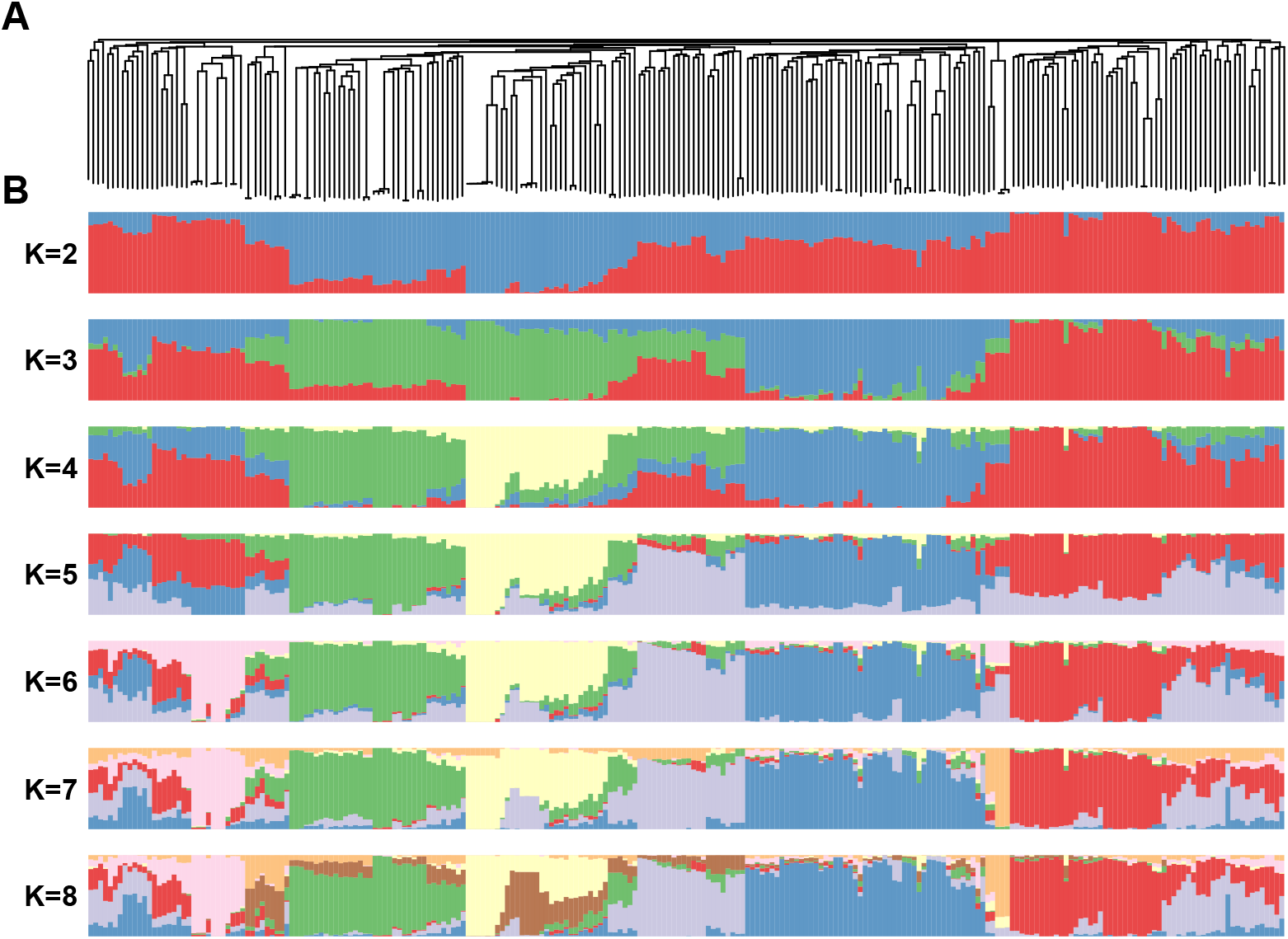
(A) Unrooted neighbor-joining (NJ) tree and (B) ancestry coefficients of 244 B1K+ accessions estimated by using *ALStructure* with *K* = *2-8*. Accessions are sorted according to the NJ tree. With *K* = *4*, red, blue, green, and yellow bars correspond to the Northern, Coastal, Eastern Desert and Southern Desert genetic clusters, respectively, as in Figure 1.

To evaluate genetic differentiation between four clusters, we computed pairwise *F_ST_* and Nei’s genetic distances. The Southern Desert cluster is most strongly isolated from the other three clusters (Table 1). With respect to the genomic pattern of differentiation, *F_ST_* values were highest in the pericentromeric regions of chromosome 2H, 3H, 4H, 5H, and 6H. The Southern Desert cluster differentiated from the other three clusters in the most of genome except the pericentromeric regions of chromosome 3H and 4H (Figure S7).

**Table 1.** *F_ST_* and Nei’s genetic distances between four genetic clusters.

| | | $F_{ST}$ | | | |
| --- | --- | --- | --- | --- | --- |
|  |  | North | Coast | Eastern Desert | Southern Desert |
| Nei's<br>Distance | North | - | 0.1124 | 0.1149 | 0.2593 |
|  | Coast | 0.0209 | - | 0.1321 | 0.2533 |
|  | Eastern Desert | 0.0216 | 0.0248 | - | 0.2125 |
|  | Southern Desert | 0.0508 | 0.0482 | 0.0389 | - |

A joint PCA of B1K+ and IPK accessions was consistent with major clusters identified in B1K+ and showed that B1K+ accessions overlapped with a large proportion of the IPK collection (Figure 3A). To visualize the genetic relationship between B1K+ accessions and IPK accessions with different origins, we selected 72 geographically distinct accessions used in a previous study (Russell et al. 2016). On the first PC axis, most of the 72 geographically diverse accessions collected from western and central Asian countries separated from B1K+ accessions but clustered more closely to the two desert clusters than to the northern and coastal clusters (Figure 3A). Because an unbalanced sample size of accessions from Israel (616 of 1,365 accessions) may bias the PC axes, we performed another joint PCA by projecting 1,293 accessions onto PC spaces of 72 geographically distinct accessions. This approach can avoid the misinterpretation of sample origin and migration based on PC (McVean 2009). The PC projection showed that accessions typically clustered by geographic origin, as reported in previous studies (Milner et al. 2019; Russell et al. 2016), and B1K+ accessions were concentrated in a small area of PC space (Figure 3B).

**Figure 3.**
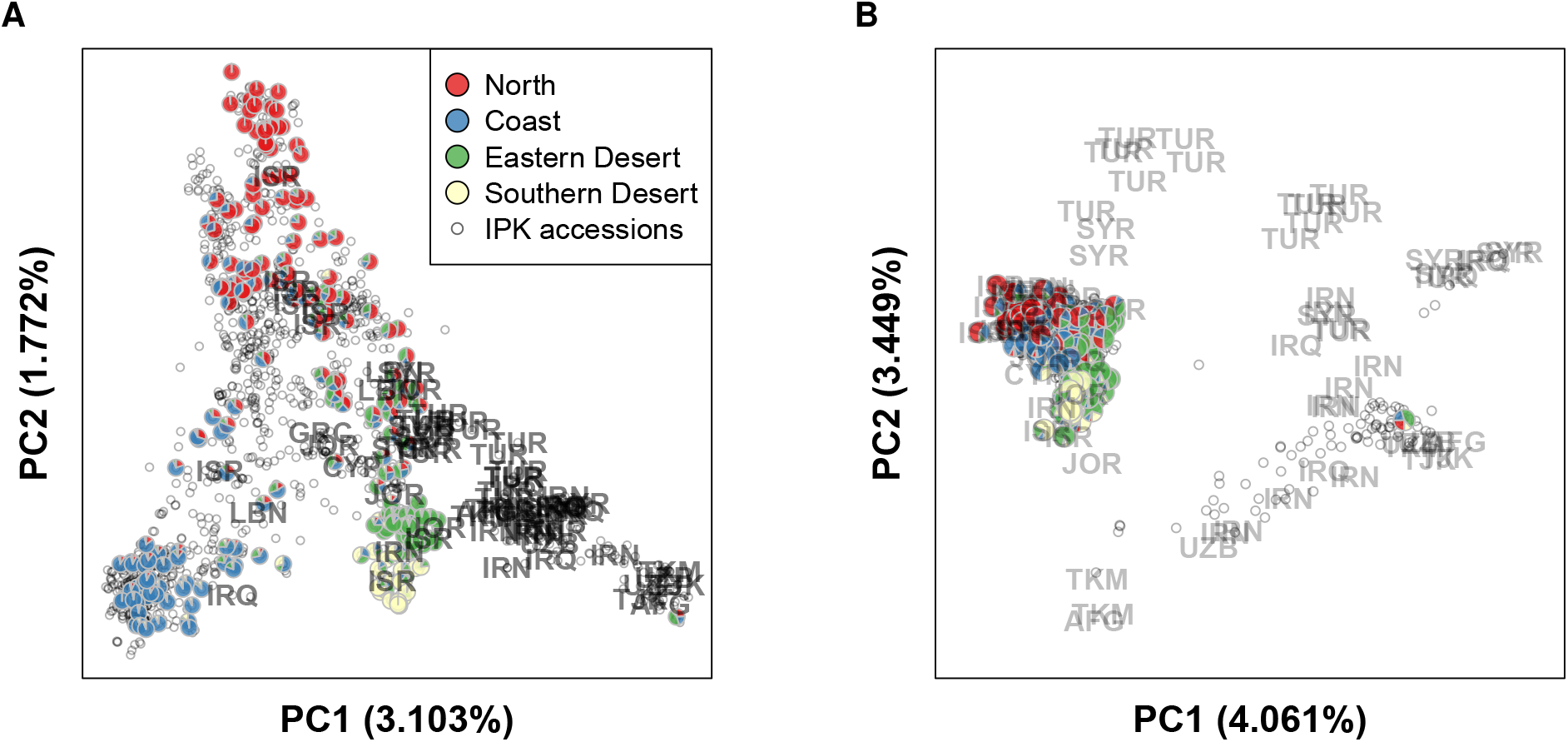
PCA plots of 244 B1K+accessions and 1,121 accessions from IPK genebank. (A) PCA performed with all of the available accessions. (B) PCA performed with 72 geographically diverse accessions and projecting the remaining accessions to PC spaces. Pie charts represent ancestry coefficients of 244 B1K+ accessions estimated by using *ALStructure* with *K*= *4*. Gray open dots represent IPK accessions.

### Geographical pattern of gene flow

To identify barriers limiting gene flow within the Levant region, we performed an analysis using *EEMS* (Petkova et al. 2016) that revealed uneven gene flow across the landscape. The area of low gene flow rates corresponds closely to geographic barriers, including the Sea of Galilee, the Jordan Valley, and the Judea and Samaria mountain ridges (Figure 1C). A Wilcoxon test supported the association between geographic barriers and lower gene flow rates (*p* < 2.2 × 10^-22^; Figure S8 A). In addition, *EEMS* analysis showed that genetic dissimilarity between demes does not have a simple linear relationship with geographic distances (Figure S8 B), indicating that isolation by distance is not sufficient to explain genetic differentiation. The *EEMS* analysis also showed that effective genetic diversity, which is the expected genetic dissimilarity of two individuals sampled from a site (Petkova et al. 2016), decreases from north to south (Figure S8 C), suggesting higher genetic diversity in the north than the south. Furthermore, we performed *unPC* (House and Hahn 2018) based on the ratio of PC-based genetic distances to geographic distances, which is more sensitive to long-distance migration than *EEMS.* The high *unPC* score comparisons supported regions of low gene flow identified by *EEMS*, particularly the majority of significant *unPC* comparisons located in the region around the Sea of Galilee in northern Israel (Figure 1D). In addition, the comparisons with low *unPC* scores suggest potentially long-distance migration events in the north-south direction (Figure 1E).

To evaluate asymmetric gene flows, we used CBI (Lundgren and Ralph 2019), which suggested unequal gene flows in a North-South direction (Figure 1F; Table S3). There is a trend for gene flow from South (region *H*) to North (region *B*) in the western region (region *H* → *E* → *B*; Figure 1F) and an opposite trend from North (region *A*) to South (region I) in the eastern region (region *A* → *D* → *F* → *I*; Figure 1F). The strongest gene flow (3.24 with the 95% credible interval of 0.51-6.32; Table S3) is observed from populations close to Jerusalem (region *H*) to the surrounding areas of Mount Carmel (region *E*), but the gene flow in the opposite direction (region *E* → *H*) is much weaker (0.77 with the 95% credible interval of 0-2.32; Table S3). Low gene flow rates of connections across geographical barriers, such as *C* ⇌ *D* and *H* ⇌ *I*, agreed with the results of *EEMS* (Figure 1C) and *unPC* (Figure 1 D and E). Also, low gene flow rates between the Negev desert (region *J*) and its adjacent regions indicated the isolation of Southern Desert accessions, consistent with high genetic differentiation suggested by the *F_ST_* values (Table 1).

### Genetic variation explained by environment and space

To quantify relative contributions of environment and space to genomic variation, simple RDA and partial RDA were performed. RDA showed that environmental variables explained 15.12% (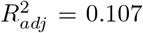; *p* = 0.0002) of SNP variation while spatial autocorrelation captured by dbMEMs, which are eigenfunctions of a spatial network (Dray et al. 2006; Legendre and Legendre 2012), explained 44.95% (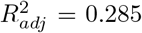; *p* = 0.0002; Figure 4A). By using partial RDA, we found 10.63% of SNP variation is jointly explained by environmental variables and spatial autocorrelation, and 4.49% (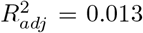; *p* = 0.0002) was exclusively explained by environmental variables (Figure 4A). Considering the confounding effect between environment and population structure, we treated ancestry coefficients (*K* = *4*) as covariates when inspecting the effect of environmental variables on SNP variation. RDA indicated that population structure explained 15.43% (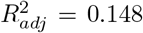; *p* = 0.0002) of SNP variation and after controlling for population structure, environmental variables exclusively explained 8.71% (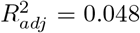; *p* = 0.0002) of SNP variation (Figure 4A).

**Figure 4.**
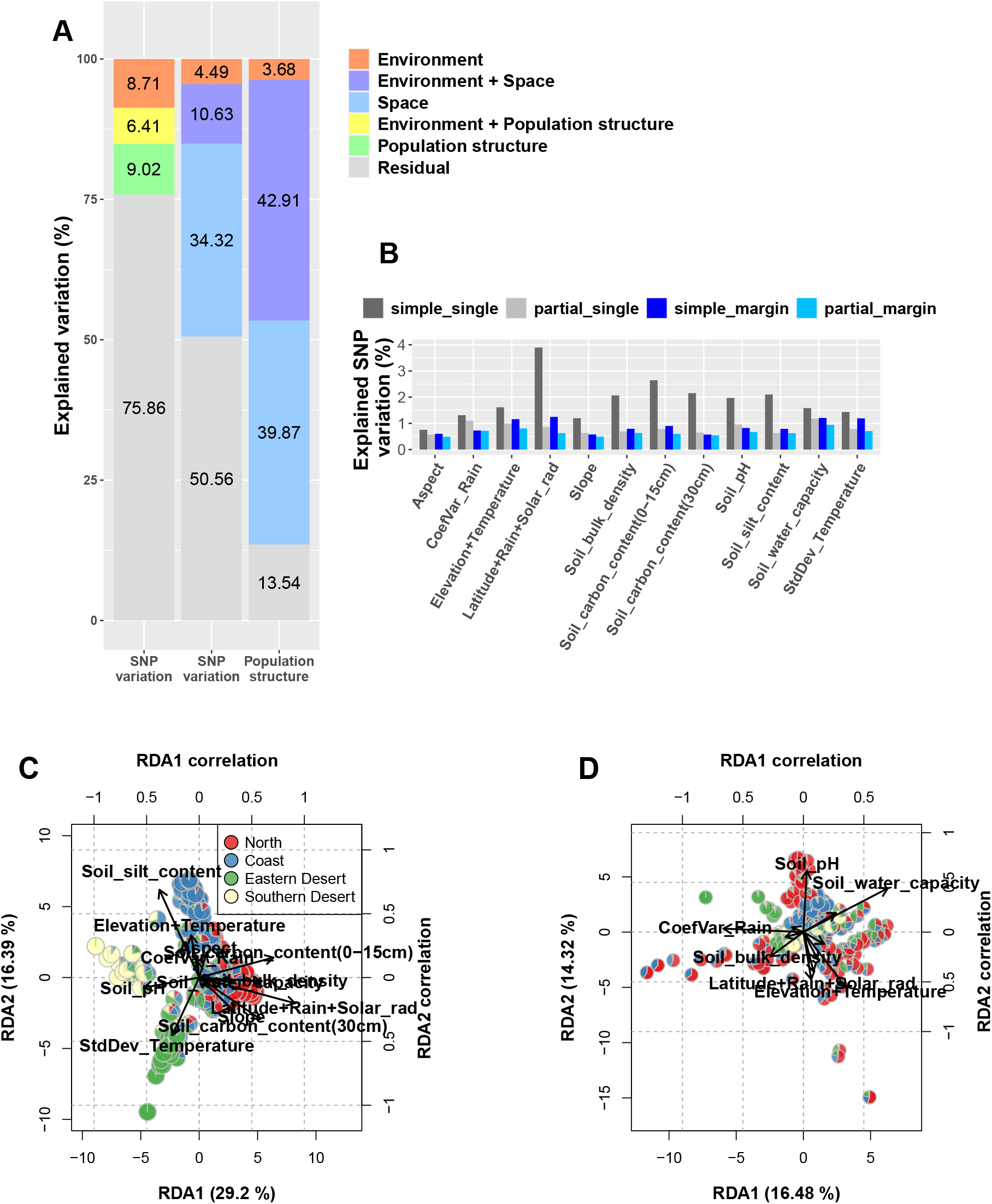
Results of variation partitioning and RDA biplots. (A) Variation partitioning of SNP variation and population structure. Left and middle columns: explained SNP variation estimated by RDA models using population structure and spatial autocorrelation as covariates, respectively. Right column: SNP variation explained by population structure. Environment, space and population structure are represented by twelve environmental variables, dbMEMs and ancestry coefficients (*K* = 4) in RDA models. (B) Percentage of SNP variation explained by environmental variables. The *simple_single* and *partial_single* show individual effects estimated based on RDA models fitting one environmental variable at a time. The *simple_margin* and *partial_margin* show marginal effects estimated based on RDA models fitting all environmental variables. The *partial_single* and *partial_margin* are estimated based on partial RDA conditioned on population structure. (C) Biplot of simple RDA. (D) Biplot of partial RDA conditioned on population structure. The arrows represent correlations of the environmental variables with RDA axes that are shown in Table S9 with details.

After confirming an association between genomic variation and the environment, we further investigated the effects of individual environmental variables. In simple RDA models separately fitting each environmental variable, permutation tests showed all of 12 environmental variables are significantly associated with SNP variation (*p* < 0.005; Table S5.1). Without constraining on population structure, the synthetic variable ‘*Latitude+Rain+Solar_rad*’ (Table S1 and S2) explained the highest proportion of SNP variation (3.89%; Figure 4 B; Table S5.1). In contrast, in RDA models controlling for population structure, ‘*Soil_water_capacity*’ explained the highest proportion of SNP variation (1.18%; Figure 4 B; Table S5.2) whereas the proportion of SNP variation explained by ‘*Latitude+Rain+Solar_rad*’ reduced to 0.86%. The variable ‘*Aspect*’ showed the lowest but significant association with SNP variation in both simple and partial RDA (Table S5.1 and S5.2). Variables ‘*Soil_water_capacity’*, ‘*CoefVar_Rain*’, which refers to coefficients of variation of precipitation of the growing season, and ‘*Aspect*’ reduced explained variation less than other environmental variables after controlling the population structure. This indicates that they are less correlated with the population structure. We also investigated marginal effects in models that incorporated all environmental variables by considering correlations between environmental variables. The variables ‘*Latitude+Rain+Solar_rad*’ and ‘*Soil_water_capacity*’ showed again the highest marginal effect in the simple RDA and partial RDA conditioned on population structure, respectively (Figure 4 B; Table S5.3 and S5.4).

RDA biplots provide further information on the relative importance of environmental gradients. The biplot of the simple RDA (Figure 4C) showed a population structure consistent with the four genetic clusters identified by *ALStructure.* The first and second RDA axes corresponded to genetic differentiation in the north-south and west-east directions, respectively (Figure 4C), and the first RDA axis is strongly (r = 0.911; Table S5) correlated with the variable ‘*Latitude+Solar_rad*’ (Figure 4C). After conditioning on spatial autocorrelation, the effects of all environmental variables decreased significantly (Figure S9 and Table S5), indicating a strong correlation of environmental gradients with spatial autocorrelation. When conditioning on population structure rather than spatial autocorrelation, two water-related variables, *“Soil_water_capacity”* (r = 0.697) and *“CoefVar_Rain”* (r = −0.662), were the most influential predictors on the first RDA axis (Figure 4D and Table S5).

With reference to our hypothesis that the diverse environments in the southern Levant are an important factor shaping populations, we quantified the relative contributions of environment and space to population structure (*K* = 4) using RDA. As expected, a high proportion of population structure explained by environmental variables (42.91 of 46.59% Figure 4A) cannot be separated from the component explained by spatial autocorrelation. Only 3.68% (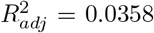; *p* = 0.0002) of population structure can be exclusively explained by environments whereas spatial autocorrelation exclusively accounted for 39.87% (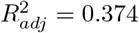; *p* = 0.0002) of population structure (Figure 4A). This result suggests that spatial autocorrelation has a larger effect on population differentiation of wild barley in the Southern Levant than environmental diversity.

### Adaptive candidates and GO enrichment

The association between genomic variation and environment motivated us to perform genome scans to identify putative adaptive loci. We first estimated LD decay to assess whether the marker density of the reduced sequenced library was sufficient to accurately identify adaptive genes in such scans. Fitting the *loess* model and Hill-Weir formula, we observed a rapid decay of LD because *r*^2^ values dropped to half of the highest values of 0.377 and 0.454 after pairwise SNP distances of 213 bp and 125 bp, respectively (Figure S10). This result indicates a possible difficulty in detecting precise locations of adaptive loci except for closely linked loci with the current marker density.

Three GEA methods, simple RDA, partial RDA, and LFMM, identified 352, 364, and 307 candidate SNPs (FDR < 0.05), respectively, and the outlier method, *BAYPASS*, identified 279 candidate SNPs (*X^T^X* > 11.05). However, candidate SNPs detected by the four methods hardly overlapped, except simple RDA and *BAYPASS* with 125 common SNPs, of which 91 are located in the pericentromeric regions of the chromosome 3H, 4H, and 5H (Figure 5; File S3). By searching 500 bp adjacent intervals of candidate SNPs, the four methods jointly identified two genes on chromosome 4H. The first gene *HORVU.MOREX.r2.4HG0308420* is close to SNPs associated with the variable ‘*Latitude+Rain+Solar_rad*’ in the LFMM analysis (File S4) and encodes an ATP-dependent RNA helicase. The second gene *HORVU.MOREX.r2.4HG0314300* is linked to SNPs associated with ‘*Elevation+Temperature*’ and which encodes a nucleolar GTP-binding protein (Figure S11; Table S6; File S4). GO term enrichment analysis identified 2 and 10 enriched GO terms based on candidate SNPs detected by simple RDA and *BAYPASS*, respectively (Table S7). No GO term was enriched based on the results of partial RDA and LFMM.

**Figure 5.**
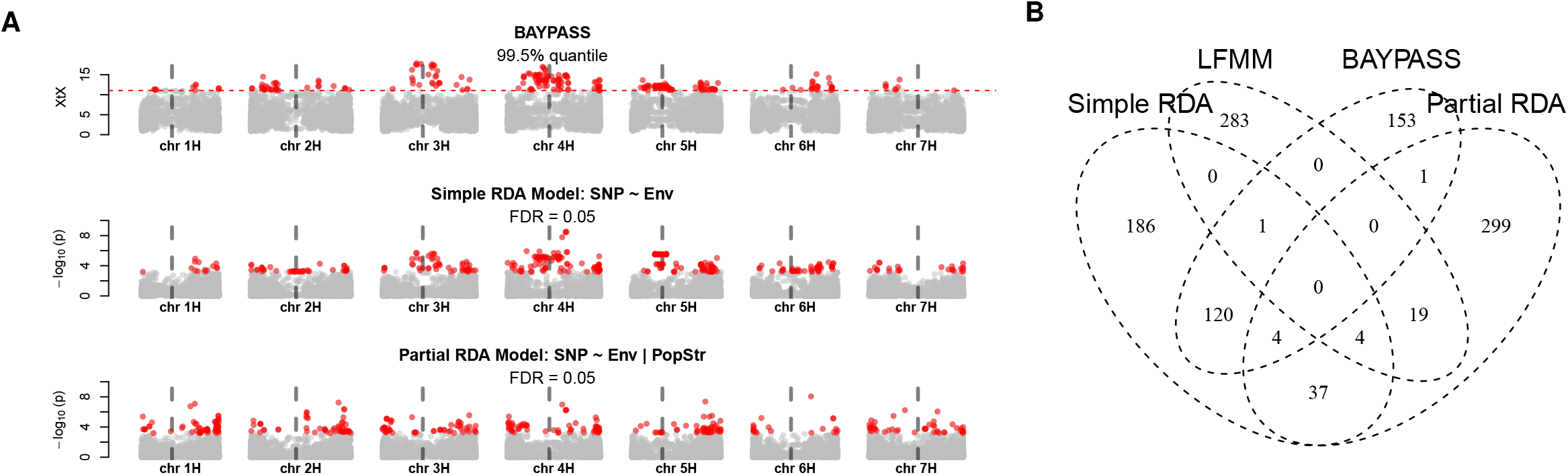
(A) Genome scans for adaptation signatures. Three manhattan plots correspond to the *BAYPASS*, simple RDA, and partial RDA conditioned on population structure. Significant SNPs are highlighted as red dots. The positions of centromeres are indicated with vertical gray dash lines. (B) Numbers of significant SNPs detected by four different methods for genome scans.

## Discussion

Our study indicated that geography and spatial autocorrelation are more predictive factors of genomic variation than environmental gradients even though the diverse environments of the Southern Levant are expected to impose strong natural selection (Hübner et al. 2009; Nevo et al. 1979). These findings imply that genomic variation of wild barley in the Southern Levant is mainly driven by neutral processes consistent with a neutralist perspective (e.g., Volis et al. 2003, 2005, 2001). However, environmental variables are still associated with a relatively small but considerable proportion of genomic variation (15.12%; Figure 4A), suggesting that natural selection and hitch-hiking may have a detectable effect on the structure of genetic diversity.

### Population structure of B1K+ and IPK genebank collection

Three clusters corresponding to eco-geographic habitats were previously characterized based on SSR markers and a SNP array developed for cultivated barley (Hübner et al. 2012) and morphological traits (Hübner et al. 2013). Our results are consistent with previous findings, except that the previously reported desert cluster (Hübner et al. 2012) split into two clusters (Figure 1 A-C), evident in the *ALStructure* analysis with *K*= *3* and *K*= *4* (Figure 2), due to not only the increased marker number but also the inclusion of additional accessions collected in the Negev Desert in 2011 (File S2). Although genetic clusters are consistent with eco-geographic habitats, the results of model-based methods should be interpreted cautiously. First, the number of ancestral populations may be overestimated due to isolation by distance (Bradburd et al. 2018). Second, the high proportion of admixed accessions (174 of 244 B1K+ accessions; 71.3%) may not result from admixture. Both spatial autocorrelation (Bradburd et al. 2018) and demographic history (Lawson et al. 2018) such as bottlenecks that likely occur in self-pollinating species (Hartfield et al. 2017), may lead to high admixture proportions in model-based methods.

The joint PCA incorporating the IPK wild barley collection indicated a strong effect of an unbalanced sample size of accessions from Israel (616 of 1,365 accessions) on a PCA (Figure 3 A and B). This comparison highlights the importance of balanced sampling in analyzing the population structure, because unequal sample sizes among groups can lead to distortion of PCs (McVean 2009). The PCA based on all accessions compressed the accessions with a broad geographical origin throughout the whole distribution range of wild barley to a cluster (Figure 3A) that did not appropriately reflect their wide geographic origin. In contrast, a PCA with a more balanced sample of accessions from the whole species range revealed that the wild barley from the Southern Levant regions comprises only a small proportion of the total diversity of wild barley (Figure 3B). However, the PCA of the complete sample revealed that accessions from Greece and Cyprus clustered with accessions from the Southern Levant (Figure 3 A and S12 A) suggesting they originated from the Southern Levant or adjacent areas without a sufficiently long history of differentiation from the ancestral populations. Likewise, 579 IPK accessions with unknown origins may be closely related to the Levant region as they strongly overlapped with our B1K+ population (Figure S12).

### Geographical pattern of gene flow

Gene flow of wild barley is expected to be limited with a low rate of outcrossing (<2%; Abdel-Ghani et al. 2004), and seed dispersal occurs mainly within 1.2 m (Volis et al. 2010). However, self-fertilizing plants can establish a population with a single seed after long-distance dispersal (Baker 1967), and the long spiky awns attached to seeds of wild barley can facilitate dispersal by zoochory. With a sufficiently long period, gene flow across landscapes may accumulate via occasional dispersal and outcrossing. *EEMS* (Petkova et al. 2016) was previously used to identify gene flow barriers in plant populations across large geographical ranges, e.g., rice (Gutaker et al. 2020) and spruce (Tsuda et al. 2016). In our data, *EEMS* revealed fine-scale patterns of gene flow attributable to geography, particularly by the Sea of Galilee and the Jordan Valley, which were not previously identified by inferring gene flow between genetic clusters (Hübner et al. 2012). These geographic separations could promote genetic differentiation within a short geographic distance that interferes with isolation-by-distance patterns (Figure S8 B).

Coalescent-based inference (CBI; Lundgren and Ralph 2019) detected trends of gene flow with opposite directions in eastern and western regions (Figure 1F), contradicting the net gene flow from north to south identified by Hübner et al. (2012). The different conclusions regarding gene flow directions in western Israel are likely due to the mappings. While Hübner et al. (2012) assigned accessions according to genetic clustering, our assignment emphasized the geographic origin. CBI gene flow rates express the probability that a population descended from another population per unit time (Lundgren and Ralph 2019), and therefore a history of recent colonization may explain gene flow trends in our data. Incorporating historical genome recombination to infer gene flow at different time periods may provide a clearer picture (Al-Asadi et al. 2019). Errors in gene flow inference can result from sampling biases, missing and erroneous genotypic values caused by low sequencing depth, and also from uneven distributions of markers due to the nature of GBS (Elshire et al. 2011; Poland et al. 2012). However, imbalanced sampling should not bias our results because *EEMS* and CBI are insensitive to unequal sample numbers (Lundgren and Ralph 2019; Petkova et al. 2016).

### Effects of environment and geographical distance on SNP variation

RDA analysis indicated that environmental gradients explained a substantial portion of SNP variation and population structure (Figure 4A). This analysis does not include all possible environmental effects because comprehensive environmental data were not available. For example, the adaptive trait *drought stress recovery* is associated with the parameter rainfall predictability in wild barley (Galkin et al. 2018), but such data are only available for some collection sites. In addition, the control for collinearity and nonlinear environmental effects that RDA does not account for may lead to unexplained genetic variation in our analysis.

Phenotypic studies suggested the importance of rainfall in the evolution of wild barley in the southern Levant (Galkin et al. 2018; Hübner et al. 2013; Volis 2011; Volis et al. 2002a,b). Our RDA analysis indicated variables related to water availability (‘*Latitude+Rain+Solar_rad*’ and ‘*Soil_water_capacity*’) as most important drivers of genomic variation (Figure 4 B-D; Table S5). It is not possible to specify the effects of individual environmental gradients because they are highly correlated. For example, we cannot separate the effect of precipitation from latitude, which is highly relevant to the flowering timing of barley (Russell et al. 2016). Unlike other environmental variables, ‘*Aspect*’ had few confounding effects with other gradients and population structure (Figure 4B). ‘*Aspect*’ was the strongest predictor when conditioned on spatial autocorrelation (Figure S9; Table S5). In the southern Levant, south-facing slopes might be more exposed to drought and heat than north-facing slopes due to higher solar radiation, resulting in significantly stronger selection within only a few hundred meters, referred to as the Evolution Canyon model (Bedada et al. 2014; Nevo et al. 2005). Our results suggest that ‘*Aspect*’ may reflect a minor but pervasive effect of microclimate in the southern Levant that cannot be represented by climate data at the current resolution. In *Mimulus guttatus*, an important locus of microgeographic adaptation was successfully identified by integrating quantitative trait loci mapping and population genomic analyses (Hendrick et al. 2016). A similar approach may be used to investigate the genetic architecture of adaptation to microclimatic conditions in wild barley.

By using dbMEMs, which model the effects of spatial autocorrelation on SNP variation, our RDA revealed that high proportions of SNP variation (45%) and population structure (83%) are explained by spatial autocorrelation (Figure 4A). The lower proportion of SNP variation explained by environments (Figure 4A) indicates that environmental selection may be an influential but not dominant driver of genetic differentiation. In contrast with our findings, environment had a significantly stronger effect than geographical distance on diversity in *Boechera stricta* (Lee and Mitchell-Olds 2011). However, in *Arabidopsis thaliana* (Lasky et al. 2012), sorghum (Lasky et al. 2015), rice (Gutaker et al. 2020) and wild tomato (Gibson and Moyle 2020), the contribution of environment was comparable and highly overlapping with geographical distance. This suggests that the isolation-by-distance is a robust and widespread pattern in a small geographic range like in our case and over large geographic scale (Gibson and Moyle 2020; Gutaker et al. 2020; Lasky et al. 2012, 2015). Complex spatial structure confounding with environmental gradients are a pervasive challenge in the study of local adaptation (de Villemereuil et al. 2014; Excoffier et al. 2009). In particular, population genetic analyses tend to be biased by spatial structure (Battey et al. 2020). For this reason, phenotypic studies using crosses between accessions and common garden experiments are also required to distinguish between genetic variation attributed to local adaptation and spatial autocorrelation. Additionally, we noted that a high percentage of SNP variation (51%; Figure 4A) remained unexplained even after incorporating dbMEMs. This could attribute to either unknown evolutionary forces independent of spatial autocorrelation or the limitation of our current dataset and methodologies.

### Lack of strong evidence to pinpoint adaptive loci

The rapid decay of LD within a few hundred base pairs (Figure S10) is consistent with similar studies of wild barley populations from the Middle East and Central Asia (Morrell et al. 2005). Reduced-representation sequencing approaches tend to have limited power in identifying adaptive loci, especially for genomes with high levels of recombination (Tiffin and Ross-Ibarra 2014). A rapid LD decay and a large genome size of ~5.3 Gb indicate that the marker density of this study may not allow precise genome scans. To account for this caveat and to control for false-positive rates, we combined the results from multiple methods of genome scans (Forester et al. 2018; Rellstab et al. 2015). Significant polymorphisms hardly overlapped between methods (Figure 5B). This observation may be explained by a (1) lack of adaptive loci with large effects, (2) strong confounding effect of population structure, and (3) limitations of the dataset. Although without strong evidence to identify adaptation genes, the genome scans based on *X^T^X* and simple RDA identified significant correlations with environmental variables and strong genetic differentiation at the pericentromeric regions of chromosome 3H, 4H, and 5H (Figure 5A). However, these associations were not observed in the partial RDA and LFMM analyses. Although the *X^T^X* statistics accounts for covariance of allele frequencies (i.e., population structure) among populations (Günther and Coop 2013), spurious signals of selection may arise if self-fertilization inflates false-positive values via strong genetic drift (Hodgins and Yeaman 2019). For this reason and because of a strong association of population structure with environments (Figure 4A), false positives are expected for the outlier and GEA methods even with a correction for population structure. The high degree of putative selection-driven differentiation is remarkable. Similar patterns of genetic differentiation in pericentromeric regions were reported in previous studies of barley (Contreras-Moreira et al. 2019; Fang et al. 2014; Wang et al. 2018), teosinte (Pyhäjärvi et al. 2013) and maize (Navarro et al. 2017). Theoretical studies suggested that adaptation with gene flow could result in divergent linkage groups of locally beneficial alleles in low recombination regions (Akerman and Bürger 2014; Bürger and Akerman 2011; Yeaman and Whitlock 2011). These conclusions were supported by simulation and empirical studies, e.g., in stickleback, sunflower, and *Arabidopsis lyrata* (Berner and Roesti 2017; Hämälä and Savolainen 2019; Renaut et al. 2013; Samuk et al. 2017). Low-recombination pericentromeric regions of wild barley were reported to have significantly higher ratios of non-synonymous to synonymous substitution (*π_a_/π_s_*) than other genomic regions (Baker et al. 2014), suggesting the tendency of accumulating genetic load in pericentromeric regions. Moreover, considering the conditional neutrality, accumulation of conditionally deleterious mutations in habitats where they are neutral can lead to genotype-environment interactions of fitness if migration is weak relative to genetic drift (Mee and Yeaman 2019). Taken together, given a weak gene flow, high rates of self-fertilization and variable recombination rates over the genome, a long-term accumulation of conditionally deleterious mutations may result in locally neutral linkage of alleles in low-recombination genomic regions creating a pattern of polymorphism that may resemble local adaptation and explain our observations in the pericentromeric regions.

### Conclusion and outlook

We observed a stronger effect of non-selective factors like geography and isolation-by-distance on total genetic diversity in the wild barley populations of the diverse and stressful environments of the Southern Levant. Nevertheless, natural selection has a small but significant influence on genomic variation, which is potentially valuable for barley breeding because water availability, i.e., precipitation and soil water capacity, was the most strongly correlated environmental variable. Outlier test and simple RDA identified genomic regions that may contribute to local adaptation, but these regions were not robustly identified by the different tests applied. One limitation of our study is therefore that only a small proportion of the wild barley genome was sequenced by the GBS approach, which is suitable for analyzing genome-wide patterns of variation and mapping of causal genes (Milner et al. 2019), but not powerful enough for pinpointing genomic targets of local adaptation. In the near future, whole genome sequencing of wild barley accessions (Sato et al. 2021) and the development of a barley pangenome (Jayakodi et al. 2020) will greatly increase the power of population genomic approaches to understand wild barley adaptation and facilitate the mining of useful alleles for plant breeding. Such approaches can be combined with common garden and transplantation experiments of wild barley genotypes to measure fitness effects in different environments (Hübner et al. 2013; Volis 2011), gene expression studies of differentially adapted genotypes (Hübner et al. 2015) and mapping populations. Given the strong influence of isolation-by-distance on genomic variation, adaptive genetic variation is likely confounded with population structure. Mapping populations with sufficient genome recombination evaluated in different environments allow for disentangling adaptive and neutral variation, as shown in such populations developed from wild and cultivated barley (Herzig et al. 2018; Wiegmann et al. 2019). Whole genome resequencing followed by computational analysis can be rationalized to analyze a large number of genotypes such as the complete B1K population. Therefore, we believe that population genomic analysis of differentially adapted crop-wild relatives will complement other approaches to understanding plant adaptation and enable use of this information for breeding.

## Acknowledgements

We are grateful to Amit Shtern for plant growth and leaf sampling, Elisabeth Kokai-Kokai for DNA extraction, and Anne Fiebig for data submission. The project at Schmid lab was supported by a Gips Schüle Foundation award “Freiräume für die Forschung” and funds of the Federal Ministry of Food and Agriculture (BMEL) based on a decision of the parliament of the Federal Republic of Germany via the Federal Office for Agriculture and Food (BLE) under the Federal Programme for Ecological Farming and Other Forms of Sustainable Agriculture (Project number 2818202615). The project at Fridman lab was supported by Israeli Science Foundation (ISF) (project number 1270/17).

## Data archiving

(The FASTQ files will be archived at European Nucleotide Archive. The geographic coordinates and environmental data is available in the File S1. The R codes will be archived at https://gitlab.com/kjschmidlab.)

