## Supplemental material for "Physical geography, isolation by distance and environmental variables shape genomic variation of wild barley (*Hordeum vulgare* L. ssp. *spontaneum*) in the Southern Levant"

**Supplementary Information**

Che-Wei Chang[1], Eyal Fridman[2], Martin Mascher[3], Axel Himmelbach[3], and Karl Schmid[1]

[1] *University of Hohenheim, Stuttgart, Germany*
[2] *Plant Sciences Institute, Agricultural Research Organization (ARO), The Volcani Center, Israel*
[3] *Leibniz Institute of Plant Genetics and Crop Plant Research (IPK), Gatersleben, Germany*

### Genotypic data filtration and processing

Six accessions are included in both IPK collection (Milner et al., 2019) and B1K collection (Hübner et al., 2009). We regarded the 6 overlapping accessions as a part of B1K collection in this study, so no accession overlaps between the 244 B1K+ accessions and 1,121 IPK's accessions in the context.

Identification of single nucleotide polymorphism (SNP) was carried out as the description of Milner et al. (2019) based on the barley 'Morex V2' assembly. The raw VCF file was first filtered by using an AWK script (bitbucket.org/ipk_dg_public/vcf_filtering) to remove loci with QUAL (mapping quality) less than 40 and genotypic calls with DP (read depth) less than 2. SNPs with a missing proportion higher than 0.2 and accessions with a missing proportion higher than 0.3 were excluded. Next, genotypic data of duplicates were merged according to the following rules: (1) retaining genotypic value if genotypes of duplicates are identical at a given SNP locus, (2) turning a given SNP locus to missing value if any genotype of duplicate conflicts with another, and (3) retaining the genotypic value if only one duplicate is not missing at a given SNP locus. Considering wild barley is predominantly self-fertilizing (Brown et al., 1978), accessions with more than 0.05 heterozygosity among all polymorphic sites were discarded, and then remaining heterozygotes in the dataset were treated as missing values. To investigate the extent of B1K+ accessions covering the genetic variation of the wild barley collection in IPK's genebank, a dataset consisting of 1,121 IPK accessions and 244 B1K+ accessions with 4,793 geographically diverse SNPs was extracted. Briefly, SNPs with a missing proportion lower than 0.2 across the whole panel and minor allele frequency (MAF) higher than 0.05 among 72 geographically diverse accessions, collected from 13 countries (Russell et al., 2016), were selected. Then, the selected SNPs were pruned by using *PLINK 1.9* (Purcell et al., 2007) to remove SNPs in linkage-disequilibrium (LD) with an $r^2$ threshold of 0.1, a window size of 50, and a step size of 5, and eventually resulted in 4,793 SNPs.

For analyses of wild barley from the southern Levant, we first extracted 58,616 SNPs with MAF higher than 0.01 and a missing proportion lower than 0.1 among 244 B1K+ accessions. For population structure analysis, LD pruning was performed to exclude redundant markers with an $r^2$ threshold of 0.1 using *PLINK 1.9* (Purcell et al., 2007) by considering the model assumption of our downstream analysis (Cabreros and Storey, 2019). LD-pruning reduced the unimputed 58,616 SNPs to 19,601 SNPs. For redundancy analysis (RDA), another dataset with no missing values was prepared because RDA requires a non-missing dataset. The whole wild barley panel, including the B1K+ collection and IPK's collection, was imputed by using *BEAGLE 5.1* (Browning et al., 2018) with the default parameters. Next, the 244 B1K+ accessions with the same 58,616

SNPs mentioned above, which have the original missing proportion lower than 0.1 among 244 accessions, were extracted. Lastly, SNPs with MAF lower than 0.05 among 244 B1K+ accessions were removed, resulting in 27,147 imputed SNPs. The imputed SNP data were coded as 0, 1, and 2 according to the counts of alternative alleles and treated as the response variable for RDA. In addition to the RDA, the imputed dataset with 27,147 SNPs was also applied to other genome scan analyses.

#### Process of generating the synthetic environmental variables and the variable selection

The climatic data including temperature, precipitation and solar radiation was downloaded from *WorldClim2* (Fick and Hijmans, 2017) with the resolution of 30 arc-seconds (~1 km). The climatic data of wild barley's growing season in Israel, typically between late October and April, was extracted to calculate the bioclimatic variables. The soil property data including a total of 11 variables of four soil layers (0cm, 5cm, 15cm, and 30cm) with the resolution of 250 m was obtained from SoilGrids (*soilgrids.org*;Hengl et al. 2017). The elevation data with the resolution of 90 m was downloaded from the SRTM database (*https://srtm.csi.cgiar.org/*). Aspects and slopes were calculated based on the elevation data using the *terrain* function in the R package *raster*. Radians of aspects were further converted to cosine values, denoting north-facing and south-facing slopes as 1 and -1, respectively. The environmental data for 244 B1K+ accessions was extracted with the function *extract* in the R package *raster* according to the geographical coordinates of their collecting locations.

To resolve the problem with collinearity, we first combined the highly correlated environmental variables to generate synthetic environmental variables. A synthetic environmental variable was defined as the first principal component scores of a group of highly correlated environmental variables. To cluster the highly correlated environmental variables that can be combined into a synthetic environmental variable, we first grouped all the environmental variables by using the complete-linkage clustering based on the dissimilarity between environmental variables. The dissimilarity of a pair of environmental variables was calculated as $1 - R^2$. The $R^2$ represents the coefficient of determination of two environmental variables. With the dendrogram of the complete-linkage clustering, we could have various clustering combinations by cutting the tree at different levels. Next, we searched the optimal clustering combinations along the dendrogram by using a customized index. The index was calculated as

$$\frac{\sum_{n=1}^{k} SS_{PC1,n}}{SS_{Total} - \sum_{n=1}^{k} SS_{PC1,n}},$$

where *k* is the number of clusters with more than one environmental variable when cutting the dendrogram at a given level. The $SS_{PC1,n}$ and $SS_{Total}$ represent the sum of squares of the first principal component calculated from $nth$ cluster of standardized environmental variables and the total sum of squares of all standardized environmental variables respectively. The $SS_{PC1,n}$ and $SS_{Total}$ were calculated by using *svd* function in R. This index was designed by regarding the sum of squares as the amount of information. Thus, the index can be interpreted as the ratio of information captured by the first PCs of grouped variables to the total information of the original data. The decrease of this index can suggest the loss of information when using the first PCs to represent the grouped variables. In other words, by maximizing the index, we could supposedly find clustering combinations producing the most informative synthetic environmental variables to represent highly correlated environmental variables. The optimal $1 - R^2$ cutoff was searched between 0 and 1 (Figure S3 A). Subsequently, the first PCs, or the synthetic environmental variables, were computed according to the optimal clustering combination (Figure S3 B). After that, the synthetic environmental variables and the non-synthetic environmental variables were further selected until variance inflation factors (VIFs) of all variables were less than 5. The selection was done by sequentially removing the environmental variable with the highest VIF, but we manually kept the variable related to the accumulated precipitation and the mean temperature ('Latitude+Rain+Solar_rad' and 'Elevation+Temperature') because the precipitation and temperature have been suggested as the most important gradients (Hübner et al., 2009). The VIFs were calculated as $\frac{1}{1 - R_j^2}$, where the $R_j^2$ is the coefficient of determination obtained by regressing variable $j$ on all the other variables (Legendre and Legendre, 2012). The procedure resulted in 12 environmental variables, including 7 synthetic environmental variables and 5 non-synthetic environmental variables (Table S1). The rotations used to generate synthetic variables are shown in Table S2.

### Classification of barrier and non-barrier pixel

To test the hypothesis that geographical barriers are significantly associated with the lower migration rate, we performed a Wilcoxon test by using migration rates estimated by *EEMS* and the geographical barriers defined according to geographical elevation. We assumed the relatively low migration areas on the landscape are due to a drastic change in elevation, such as valleys and mountains, so we identified geographical barriers by selecting the map pixels with elevation deviating from the majority of pixels (Figure S4). Specifically, we selected the pixels with the top 25%, a cut-off where pixel counts drastically decreased (Figure S4 A), of the absolute values of the standardized elevation and treated selected pixels as the geographical barriers (Figure S4 B). Thereby, pixels of the geographical map were classified as barrier pixels and non-barrier pixels, and the corresponding migration rate of each pixel was extracted from the result of *EEMS*. Finally, a Wilcoxon test was carried out to test if the migration rates are lower at the barrier pixels than the non-barrier pixels.

#### Supplementary Tables

**Table S1** Description of environmental variables used in redundancy analysis (RDA) and genome-environment association (GEA) analysis

| Environmental variable | Description | Proportion of explained variance[a] |
|---|---|---|
| Aspect | Geographical aspect | - |
| CoefVar_Rain | Coefficient of variation of precipitation of growing season | - |
| Elevation+Temperature | First PC of elevation, mean temperature, maximal temperature, and minimal temperature of growing season | 0.931 |
| Latitude+Rain+Solar_rad | First PC of latitude, accumulated precipitation, and mean solar radiation of growing season | 0.910 |
| Slope | Geographical slope | - |
| Soil_bulk_density | First PC of soil bulk density of the depths of 0 cm and 5 cm | 0.947 |
| Soil_carbon_content(0-15cm) | First PC of soil organic carbon content of the depth of 0 cm, 5 cm, and 15 cm | 0.883 |
| Soil_carbon_content(30cm) | Soil organic carbon content of the depth of 30 cm | - |
| Soil_pH | First PC of soil pH of the depth of 0 cm, 5 cm, 15 cm, and 30 cm | 0.969 |
| Soil_silt_content | First PC of silt content (2–50 micro meter) mass fraction in percentage of the depth of 0 cm, 5 cm, 15 cm, and 30 cm | 0.970 |
| Soil_water_capacity | First PC of soil water capacity with with FC = pF 2.0, 2.3, and 2.5 until wilt point of the depth of 0 cm | 0.962 |
| StdDev_Temperature | Standard deviation of temperature of growing season | - |

[a]Proportion of variance explained by the first principal component (PC) of the environmental variables used to generate the synthetic environmental variable. The rotations used to produce first PCs are shown in Table S2.

**Table S2** Rotations of first principal component (PC1) used to generate the synthetic environmental variables

| Environmental Variable | Rotation of PC1 |
|---|---|
| Aspect | - |
| CoefVar_Rain | - |
| Elevation+Temperature | $0.508 \times Elevation - 0.513 \times avg\_temp - 0.491 \times max\_temp - 0.487 \times min\_temp$ |
| Latitude+Rain+Solar_rad | $0.588 \times Latitude + 0.557 \times sum\_prec - 0.587 \times avg\_srad$ |
| Slope | - |
| Soil_bulk_density | $-0.707 \times sbd(0cm) - 0.707 \times sbd(5cm)$ |
| Soil_carbon_content(0-15cm) | $0.557 \times socc(0cm) + 0.590 \times socc(5cm) + 0.585 \times socc(15cm)$ |
| Soil_carbon_content(30cm) | - |
| Soil_pH | $0.495 \times pH(0cm) + 0.502 \times pH(5cm) + 0.503 \times pH(15cm) + 0.500 \times pH(30cm)$ |
| Soil_silt_content | $-0.499 \times sltc(0cm) - 0.502 \times sltc(5cm) - 0.503 \times sltc(15cm) - 0.495 \times sltc(30cm)$ |
| Soil_water_capacity | $-0.575 \times swc(pF2.0; 0cm) - 0.582 \times \times swc(pF2.3; 0cm) - 0.575 \times \times swc(pF2.5; 0cm)$ |
| StdDev_Temperature | - |

$sum\_prec$: Accumulated precipitation of growing season
$avg\_srad$: Average solar radiation of growing season
$avg\_temp$: Average temperature of growing season
$max\_temp$: Maximum temperature of growing season
$min\_temp$: Minimum temperature of growing season
$bd$: Soil bulk density (depth of soil layer is shown in the parenthesis)
$swc$: Soil water capacity (field capacity, referred to pF, and depth of soil layer are shown in the parenthesis)
$socc$: Soil organic carbon content (depth of soil layer is shown in the parenthesis)
$pH$: Soil pH value (depth of soil layer is shown in the parenthesis)
$sltc$: Silt content (depth of soil layer is shown in the parenthesis)

**Table S3** Gene flow rate and coalescence rate of 10 geographical clusters inferred by coalescence-based method. The rows and columns respectively indicate the source and sink of gene flow. The coalescence rates are shown in diagonal. The 95% credible intervals are shown in the parentheses.

| | To | | | | | | | | | |
|---|---|---|---|---|---|---|---|---|---|---|
| From | A | B | C | D | E | F | G | H | I | J |
| A | 0.9304 (0-2.04) | 1.1481 (0-2.99) | 1.3852 (0-4.46) | 2.6973 (1.1-4.69) | - | - | - | - | - | - |
| B | 0.66 (0-2.27) | 1.2995 (0.42-2.81) | 0.7353 (0-2.73) | - | 0.7096 (0-2.15) | 0.2949 (0-0.89) | - | - | - | - |
| C | 1.2967 (0-3.99) | 0.7587 (0-2.65) | 0.9776 (0.28-1.96) | 0.5604 (0-1.71) | 0.3461 (0-1.29) | 0.5628 (0-1.29) | - | - | - | - |
| D | 0.4697 (0-1.84) | - | 0.4442 (0-1.61) | 1.3294 (0.42-2.64) | 0.4024 (0-1.62) | 2.9335 (1.49-4.88) | - | - | - | - |
| E | - | 1.4597 (0-3.5) | 0.3548 (0-1.42) | 0.3587 (0-1.32) | 0.4086 (0-1) | 0.8163 (0-1.91) | 0.5671 (0-1.81) | 0.7746 (0-2.32) | 0.5595 (0-1.71) | - |
| F | - | 0.534 (0-1.6) | 0.4686 (0-1.59) | 0.6024 (0-2.35) | 0.4509 (0-1.89) | 2.3576 (0.88-4.33) | 0.2834 (0-0.83) | 0.5712 (0-1.8) | 2.0077 (0.84-3.55) | - |
| G | - | - | - | - | 0.4679 (0-1.39) | 0.1673 (0-0.56) | 1.2041 (0.4-2.2) | 1.1336 (0-3.12) | - | 0.0865 (0-0.36) |
| H | - | - | - | - | 3.2381 (0.51-6.32) | 1.1729 (0-2.36) | 2.0253 (0-3.92) | 1.7529 (0.61-3.27) | 0.0983 (0-0.41) | 0.0655 (0-0.3) |
| I | - | - | - | - | 0.154 (0-0.54) | 0.242 (0-0.8) | - | 0.1216 (0-0.4) | 0.5207 (0.18-1.14) | 0.253 (0-1.16) |
| J | - | - | - | - | - | - | 0.1044 (0-0.46) | 0.5311 (0.04-0.91) | 0.538 (0-0.99) | 0.6713 (0.2-1.19) |

**Table S4** Effect of environmental variables estimated by RDA models. Environmental variables are ordered according to the explained variation. P-values are computed by permutation tests with 5,000 iterations.

| Table S5.1 | Environmental variable | F | Var | Var (%) | p-value |
|---|---|---|---|---|---|
| Individual effect of environmental variables | Latitude+Rain+Solar_rad | 9.8046 | 553.6372 | 3.8937 | 0.0002 |
| | Soil_carbon_content(0-15cm) | 6.5739 | 376.0355 | 2.6447 | 0.0002 |
| | Soil_carbon_content(30cm) | 5.3230 | 306.0195 | 2.1522 | 0.0002 |
| | Soil_silt_content | 5.1947 | 298.7987 | 2.1015 | 0.0002 |
| | Soil_bulk_density | 5.1081 | 293.9198 | 2.0671 | 0.0002 |
| | Soil_pH | 4.8677 | 280.3609 | 1.9718 | 0.0002 |
| | Elevation+Temperature | 3.9666 | 229.2979 | 1.6127 | 0.0002 |
| | Soil_water_capacity | 3.8998 | 225.4961 | 1.5859 | 0.0002 |
| | StdDev_Temperature | 3.5253 | 204.1527 | 1.4358 | 0.0002 |
| | CoefVar_Rain | 3.2164 | 186.4980 | 1.3116 | 0.0002 |
| | Slope | 2.9324 | 170.2299 | 1.1972 | 0.0002 |
| | Aspect | 1.8359 | 107.0534 | 0.7529 | 0.0002 |

| Table S5.2 | Environmental variable | F | Var | Var (%) | p-value |
|---|---|---|---|---|---|
| Individual effect of environmental variables conditioned on population structure | Soil_water_capacity | 3.3934 | 168.3390 | 1.1839 | 0.0002 |
| | CoefVar_Rain | 3.1455 | 156.2026 | 1.0986 | 0.0002 |
| | Elevation+Temperature | 2.8432 | 141.3632 | 0.9942 | 0.0002 |
| | Soil_pH | 2.7389 | 136.2396 | 0.9582 | 0.0002 |
| | Latitude+Rain+Solar_rad | 2.4531 | 122.1676 | 0.8592 | 0.0002 |
| | StdDev_Temperature | 2.2437 | 111.8356 | 0.7865 | 0.0002 |
| | Soil_carbon_content(0-15cm) | 2.2237 | 110.8457 | 0.7796 | 0.0002 |
| | Soil_bulk_density | 1.9666 | 98.1368 | 0.6902 | 0.0002 |
| | Soil_carbon_content(30cm) | 1.8504 | 92.3804 | 0.6497 | 0.0002 |
| | Slope | 1.8143 | 90.5932 | 0.6371 | 0.0002 |
| | Soil_silt_content | 1.7954 | 89.6590 | 0.6306 | 0.0002 |
| | Aspect | 1.6250 | 81.2028 | 0.5711 | 0.0002 |

| Table S5.3 | Environmental variable | F | Var | Var (%) | p-value |
|---|---|---|---|---|---|
| Marginal effect of environmental variables | Latitude+Rain+Solar_rad | 3.3906 | 177.1453 | 1.2459 | 0.0002 |
| | Soil_water_capacity | 3.2845 | 171.6034 | 1.2069 | 0.0002 |
| | StdDev_Temperature | 3.2423 | 169.3995 | 1.1914 | 0.0002 |
| | Elevation+Temperature | 3.1535 | 164.7580 | 1.1587 | 0.0002 |
| | Soil_carbon_content(0-15cm) | 2.4493 | 127.9675 | 0.9000 | 0.0002 |
| | Soil_pH | 2.2410 | 117.0829 | 0.8234 | 0.0002 |
| | Soil_bulk_density | 2.1565 | 112.6705 | 0.7924 | 0.0002 |
| | Soil_silt_content | 2.1446 | 112.0493 | 0.7880 | 0.0004 |
| | CoefVar_Rain | 1.9776 | 103.3233 | 0.7267 | 0.0002 |
| | Aspect | 1.6364 | 85.4935 | 0.6013 | 0.0018 |
| | Slope | 1.5674 | 81.8895 | 0.5759 | 0.0024 |
| | Soil_carbon_content(30cm) | 1.5571 | 81.3511 | 0.5721 | 0.0054 |

| Table S5.4 | Environmental variable | F | Var | Var (%) | p-value |
|---|---|---|---|---|---|
| Marginal effect of environmental variables conditioned on population structure | Soil_water_capacity | 2.8405 | 134.3772 | 0.9451 | 0.0002 |
| | Elevation+Temperature | 2.4234 | 114.6438 | 0.8063 | 0.0002 |
| | CoefVar_Rain | 2.1510 | 101.7564 | 0.7157 | 0.0002 |
| | StdDev_Temperature | 2.1193 | 100.2602 | 0.7051 | 0.0002 |
| | Soil_pH | 2.0162 | 95.3822 | 0.6708 | 0.0006 |
| | Soil_bulk_density | 1.8989 | 89.8333 | 0.6318 | 0.0006 |
| | Latitude+Rain+Solar_rad | 1.8864 | 89.2404 | 0.6276 | 0.0002 |
| | Soil_silt_content | 1.8743 | 88.6696 | 0.6236 | 0.0004 |
| | Soil_carbon_content(0-15cm) | 1.7986 | 85.0871 | 0.5984 | 0.0006 |
| | Soil_carbon_content(30cm) | 1.6314 | 77.1770 | 0.5428 | 0.0034 |
| | Slope | 1.4805 | 70.0406 | 0.4926 | 0.0080 |
| | Aspect | 1.4777 | 69.9055 | 0.4916 | 0.0090 |

**Table S5** Correlation of environmental variables with fitted site scores on first four RDA axes. The correlations were evaluated based on the RDA models (1) without covariates (Simple RDA), (2) conditioned on spatial autocorrelation with dbMEMs (Partial RDA dbMEM), and (3) conditioned on population structure with ancestry coefficients (Partial RDA PopStr)

| | Simple RDA | | | | Partial RDA dbMEM | | | | Partial RDA PopStr | | | |
|---|---|---|---|---|---|---|---|---|---|---|---|---|
| | RDA1 | RDA2 | RDA3 | RDA4 | RDA1 | RDA2 | RDA3 | RDA4 | RDA1 | RDA2 | RDA3 | RDA4 |
| Aspect | 0.067 | 0.185 | 0.251 | 0.3 | 0.274 | -0.365 | -0.121 | 0.237 | 0.269 | 0.193 | 0.072 | 0.038 |
| CoefVar_Rain | -0.015 | 0.126 | 0.543 | -0.604 | 0.013 | 0.018 | 0.017 | 0.008 | -0.662 | 0.035 | 0.106 | 0.584 |
| Elevation+Temperature | -0.077 | 0.33 | -0.751 | -0.193 | 0.001 | -0.011 | 0.008 | 0.004 | 0.353 | -0.557 | 0.265 | -0.046 |
| Latitude+Rain+Solar_rad | 0.911 | -0.201 | 0.023 | -0.197 | -0.004 | -0.007 | 0.005 | 0.002 | 0.065 | -0.475 | -0.049 | 0.027 |
| Slope | 0.335 | -0.264 | -0.027 | -0.178 | 0.038 | -0.065 | 0.106 | -0.069 | -0.146 | -0.049 | 0.371 | -0.198 |
| Soil_bulk_density | 0.59 | -0.03 | 0.037 | -0.44 | 0.041 | -0.036 | 0.021 | 0.044 | -0.272 | -0.243 | 0.176 | -0.045 |
| Soil_carbon_content(0-15cm) | 0.706 | 0.157 | 0.153 | -0.218 | -0.019 | 0.022 | -0.020 | -0.001 | 0.084 | -0.396 | 0.107 | -0.21 |
| Soil_carbon_content(30cm) | 0.582 | -0.325 | 0.209 | 0.221 | 0.086 | 0.044 | -0.015 | 0.029 | 0.169 | -0.118 | -0.23 | -0.254 |
| Soil_pH | -0.526 | -0.078 | -0.224 | 0.326 | 0.021 | 0.077 | 0.025 | 0.079 | 0.028 | 0.615 | 0.354 | -0.144 |
| Soil_silt_content | -0.377 | 0.687 | 0.203 | -0.238 | 0.149 | -0.095 | 0.027 | -0.078 | -0.093 | 0.049 | 0.12 | 0.382 |
| Soil_water_capacity | 0.325 | -0.039 | -0.254 | 0.845 | -0.016 | -0.004 | -0.042 | -0.053 | 0.697 | 0.44 | -0.18 | -0.072 |
| StdDev_Temperature | -0.257 | -0.453 | -0.113 | 0.216 | -0.020 | -0.038 | -0.006 | -0.003 | 0.118 | -0.272 | -0.544 | -0.267 |

**Table S6** Genes within 500 bp upstream or downstream of the candidate SNPs jointly detected by four methods

| Gene | Chr. | Start (bp) | End (bp) | Position of candidate SNP (bp) | Method | Environmental variable | Gene annotation |
|---|---|---|---|---|---|---|---|
| HORVU.MOREX.r2.4HG0308420 | chr4H | 312,111,636 | 312,189,673 | 312,146,232 | Simple RDA ($p$ = 2.007e-06; $q$ = 0.002) | - | ATP-dependent RNA helicase |
| | | | | 312,146,232 | Partial RDA ($p$ = 2.641e-04; $q$ = 0.031) | - | |
| | | | | 312,146,215 | LFMM ($p$ = 1.940e-06; $q$ = 0.028) | Latitude+Rain+Solar_rad | |
| | | | | 312,146,232 | BAYPASS ($X^T X$ = 11.303) | - | |
| HORVU.MOREX.r2.4HG0314300 | chr4H | 392,019,895 | 392,026,683 | 392,022,112 | Simple RDA ($p$ = 3.183e-09; $q$ = 3.245e-05) | - | Nucleolar GTP-binding protein 2 |
| | | | | 392,022,112 | Partial RDA ($p$ = 5.620e-07; $q$ = 0.001) | - | |
| | | | | 392,021,665 | LFMM ($p$ = 1.372e-4; $q$ = 0.0364) | Elevation+Temperature | |
| | | | | 392,021,634 | BAYPASS ($X^T X$ = 14.690) | - | |

**Table S7** Result of gene ontology (GO) term enrichment

| | Enriched GO | P-value | FDR | GO definition |
|---|---|---|---|---|
| Simple RDA | GO:0004556 | 2.7640e-07 | 0.0014 | Catalysis of the endohydrolysis of (1->4)-alpha-D-glucosidic linkages in polysaccharides containing three or more alpha-(1->4)-linked D-glucose units. |
| | GO:0103025 | 2.3737e-07 | 0.0014 | Catalysis of the reaction: n H2O + a 1,4-alpha-D-glucan = alpha-maltohexaose + a 1,4-alpha-D-glucan |
| *BAYPASS* | GO:0006631 | 8.2636e-06 | 0.0077 | The chemical reactions and pathways involving fatty acids, aliphatic monocarboxylic acids liberated from naturally occurring fats and oils by hydrolysis. |
| | GO:0009735 | 1.1465e-06 | 0.0029 | Any process that results in a change in state or activity of a cell or an organism (in terms of movement, secretion, enzyme production, gene expression, etc.) as a result of a cytokinin stimulus. |
| | GO:0009236 | 2.5754e-05 | 0.0203 | The chemical reactions and pathways resulting in the formation of cobalamin (vitamin B12), a water-soluble vitamin characterized by possession of a corrin nucleus containing a cobalt atom. |
| | GO:0090351 | 1.18776e-07 | 0.0004 | The process whose specific outcome is the progression of the seedling over time, beginning with seed germination and ending when the first adult leaves emerge. |
| | GO:0006635 | 2.4051e-06 | 0.0038 | A fatty acid oxidation process that results in the complete oxidation of a long-chain fatty acid. Fatty acid beta-oxidation begins with the addition of coenzyme A to a fatty acid, and occurs by successive cycles of reactions during each of which the fatty acid is shortened by a two-carbon fragment removed as acetyl coenzyme A; the cycle continues until only two or three carbons remain (as acetyl-CoA or propionyl-CoA respectively). |
| | GO:0009062 | 5.8859e-06 | 0.0063 | The chemical reactions and pathways resulting in the breakdown of a fatty acid, any of the aliphatic monocarboxylic acids that can be liberated by hydrolysis from naturally occurring fats and oils. Fatty acids are predominantly straight-chain acids of 4 to 24 carbon atoms, which may be saturated or unsaturated; branched fatty acids and hydroxy fatty acids also occur, and very long chain acids of over 30 carbons are found in waxes. |
| | GO:0009845 | 1.1054e-07 | 0.0004 | The physiological and developmental changes that occur in a seed commencing with water uptake (imbibition) and terminating with the elongation of the embryonic axis. |
| | GO:0004325 | 2.4976e-06 | 0.0038 | Catalysis of the reaction: protoheme = Fe(2+) + protoporphyrin IX. |
| | GO:0046487 | 1.1351e-09 | 1.1649e-05 | The chemical reactions and pathways involving glyoxylate, the anion of glyoxylic acid, HOC-COOH. |
| | GO:0019395 | 1.7175e-05 | 0.0147 | The removal of one or more electrons from a fatty acid, with or without the concomitant removal of a proton or protons, by reaction with an electron-accepting substance, by addition of oxygen or by removal of hydrogen. |

#### Supplementary Figures

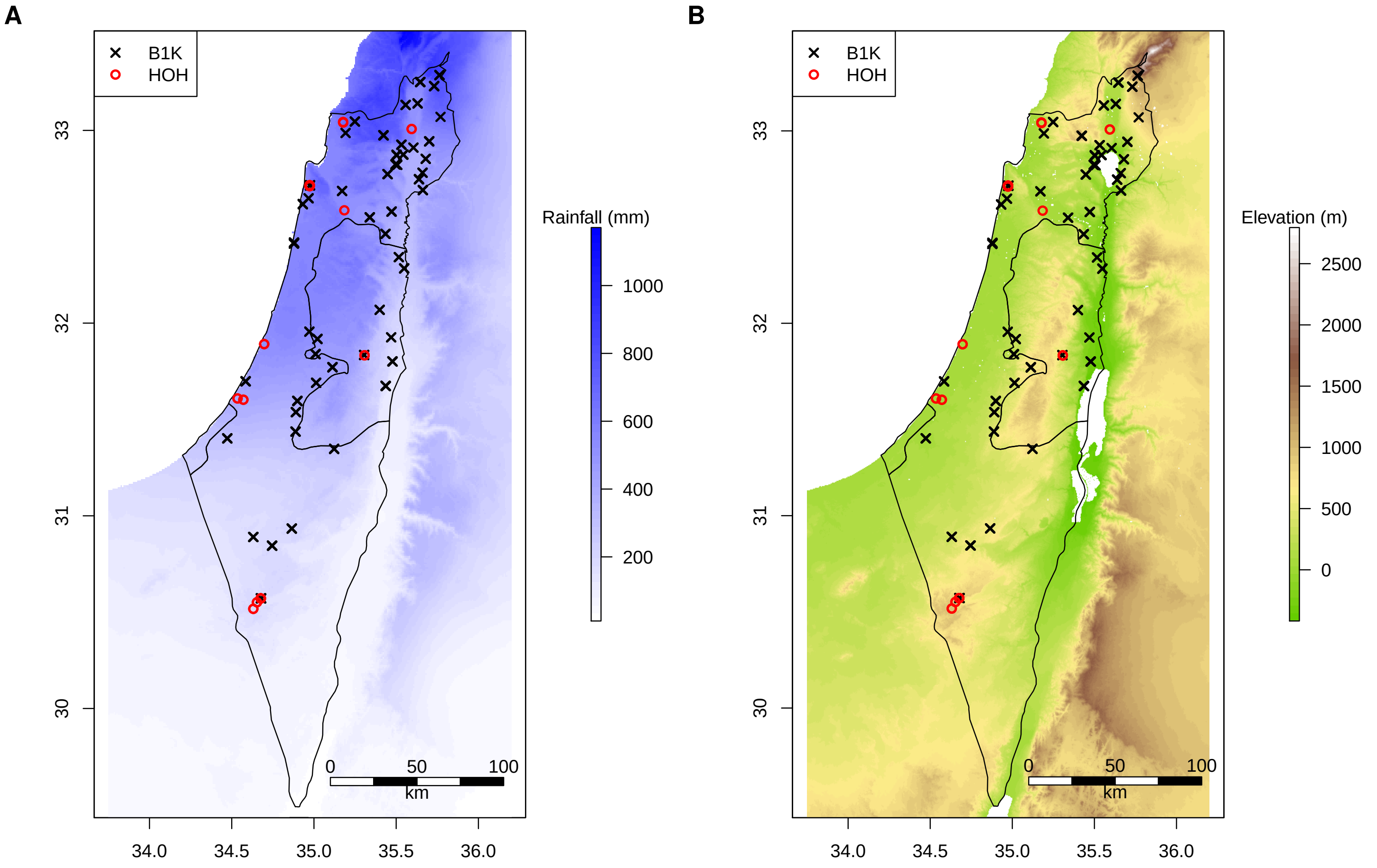


**Figure S1** Collection sites of 244 B1K+ accessions. (A) Map with the accumulated rainfall between October and April. (B) Map with the geographical elevation. The black crosses and red circles represent the collection sites of B1K accessions and unpublished accessions stored at the University of Hohenheim (HOH collection), respectively

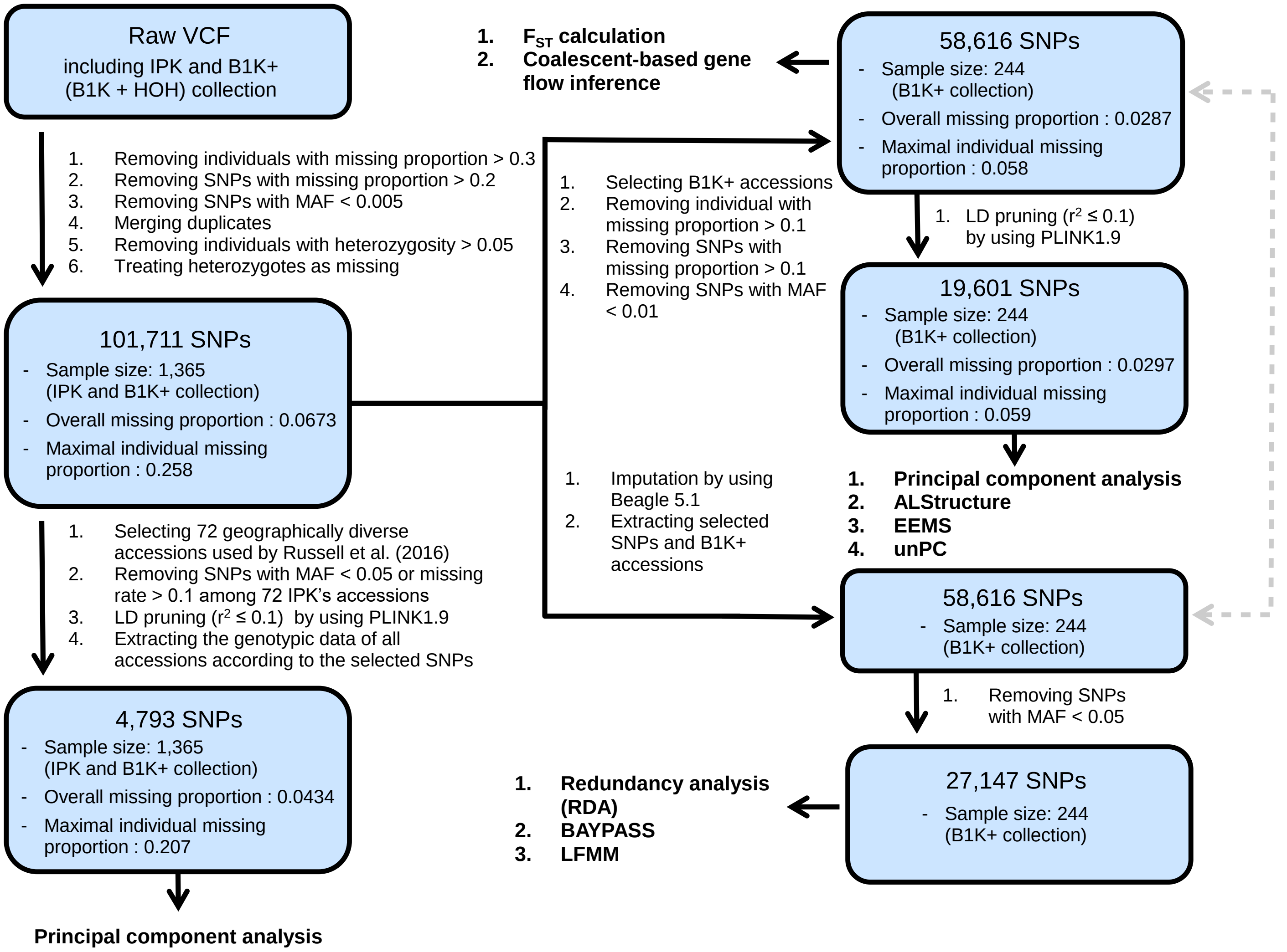


**Figure S2** Workflow of genotypic data filtration. The double arrow with the gray dash line represents that two datasets are consisting of the same 58,616 SNP loci.

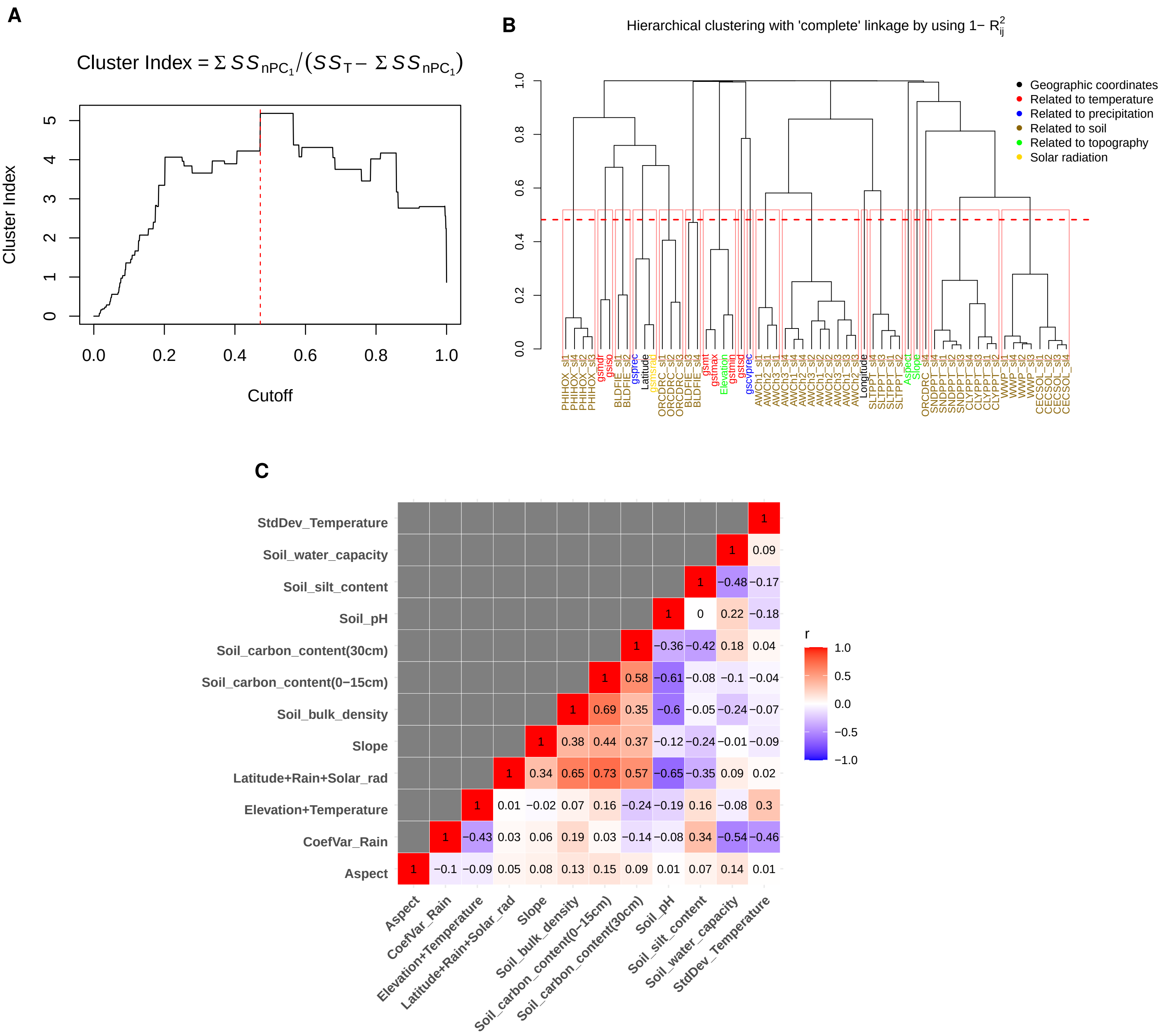


**Figure S3** Procedure of generating synthetic environmental variables. (A) Relationship of cutoff for clustering and corresponding cluster index. The y-axis represents the customized cluster index, and the x-axis represents $1 - R^2$ that is used as a cutoff for the dendrogram. The vertical red dash line shows the optimal cutoff, which is 0.472 (B) Dendrogram of hierarchical clustering based on the $1 - R^2$. The horizontal red dash line shows the optimal cutoff used to determine the clustering combinations for generating synthetic environmental variables. (C) Correlation between the retained 12 environmental variables.

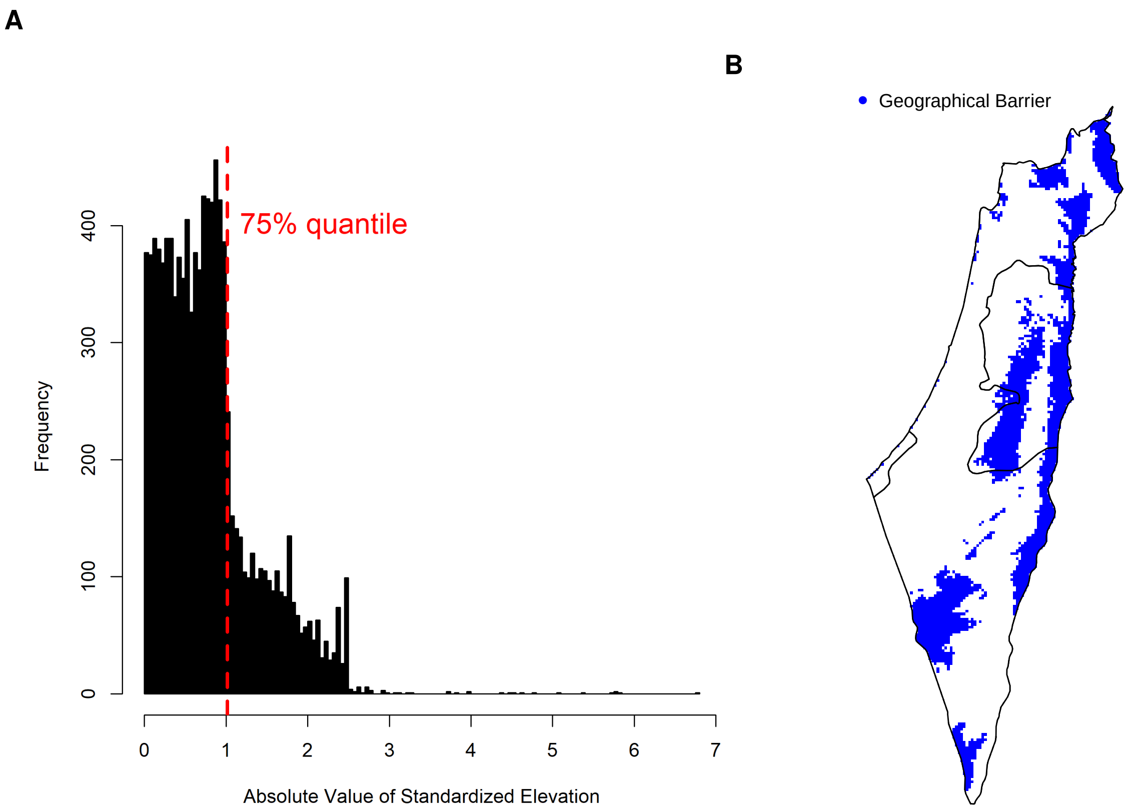


**Figure S4** Classification of barrier and non-barrier pixels. (A) Histogram of absolute value of standardized elevation. The map pixels with elevation deviating from the majority of pixels are defined as geographical barriers. The top 25% cutoff is determined by the drastic decrease of pixel counts. (B) Geographical distribution of barrier pixels.

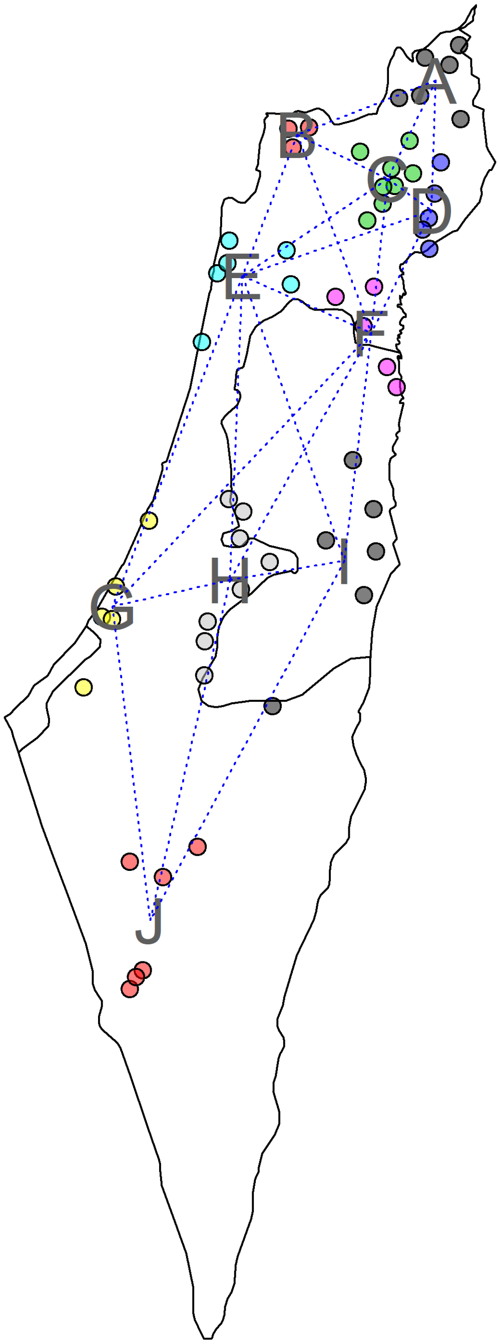


**Figure S5** Geographical clusters and network used for the coalescent-based gene flow inference. The blue dash lines represent the edges of network which allows the movement between the connected geographical regions in the gene flow model.

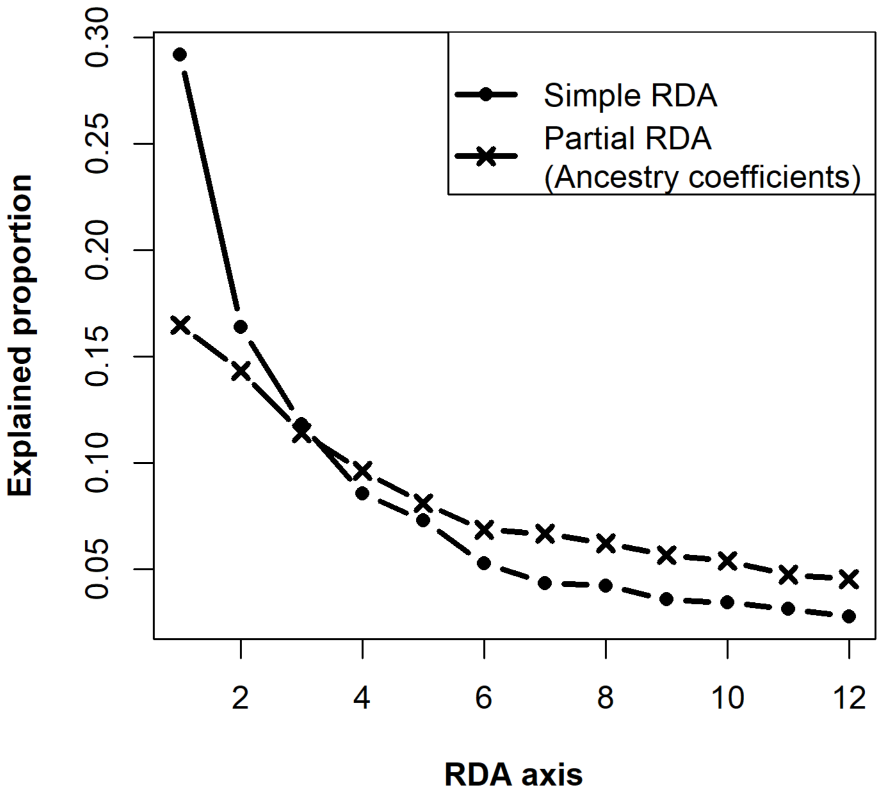


**Figure S6** Explained proportion of variation of RDA axes. The lines with dots and crosses show the result of simple RDA and partial RDA conditioned on population structure, respectively. The first four RDA axes are selected to compute Mahalanobis distances.

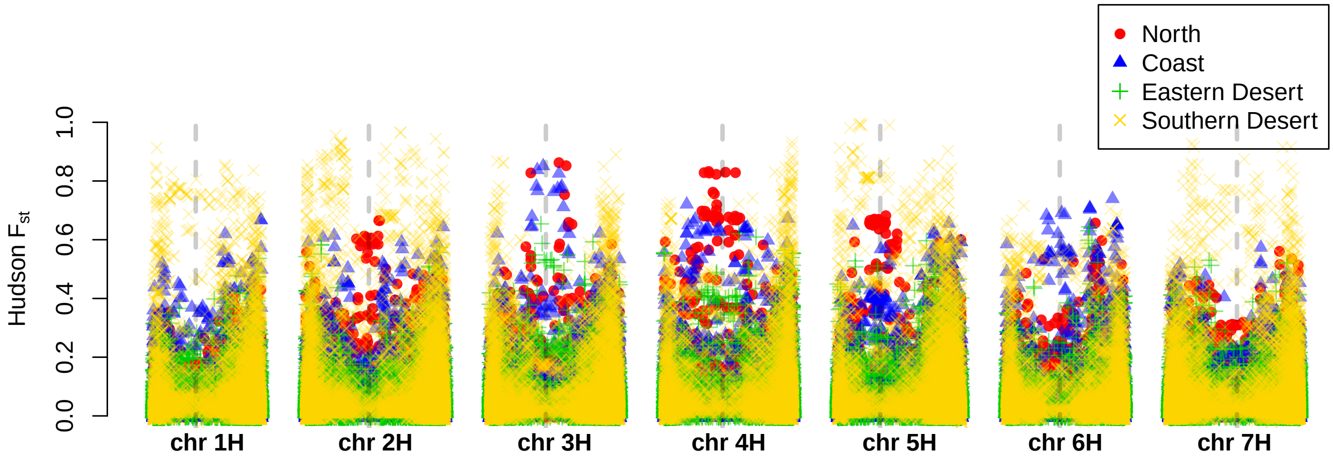


**Figure S7** $F_{ST}$ values along genome. The position of centromeres is shown by the vertical gray dash lines. The dots represent $F_{ST}$ calculated between a specific genetic cluster and three remaining genetic clusters as a whole.

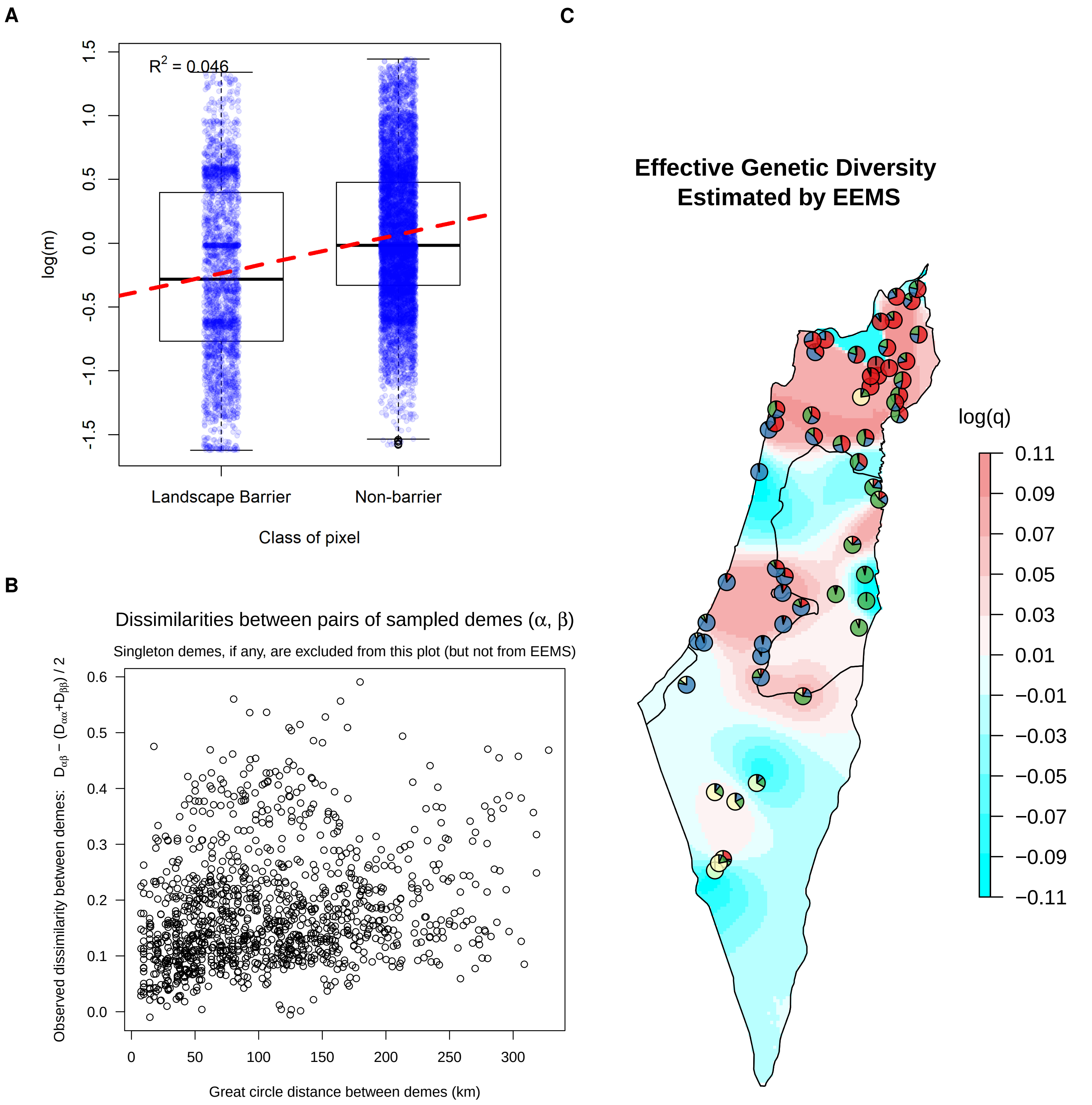


**Figure S8** Result of EEMS. (A) Logarithm of gene flow rate ($log[m]$) estimated by EEMS classified according to barrier and non-barrier pixels. (B) Relationship between observed genetic dissimilarity computed by EEMS and geographical distances. (C) Effective genetic diversity surface estimated by EEMS. Effective genetic diversity ($q$) is the expected genetic dissimilarity of two individuals sampled from a location. The pie charts represent the average ancestry coefficients of accessions in each deme.

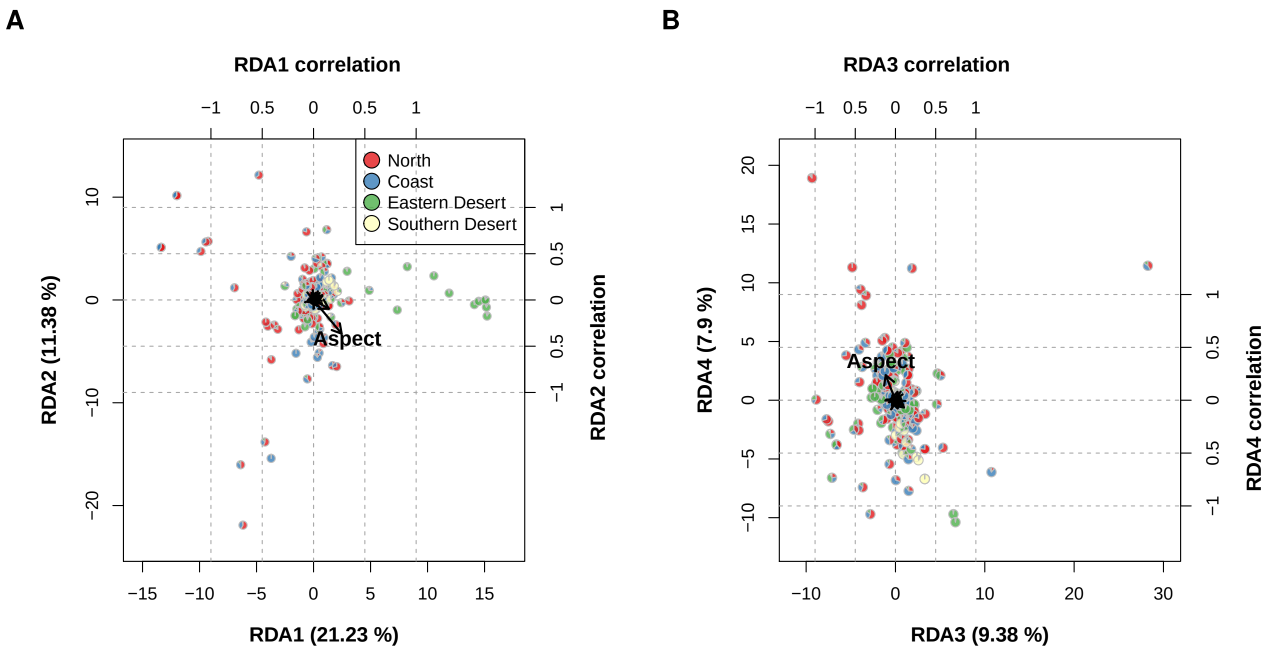


**Figure S9** Biplot of RDA conditioned on spatial autocorrelation. (A) Biplot of the first and second RDA axes. (B) Biplot of the third and fourth RDA axes. The arrows represent correlations of the environmental variables with RDA axes that are shown in Table S9 in details. The pies represent ancestry coefficients of individuals. The coordinates of pies correspond to site scores of individuals on RDA axes.

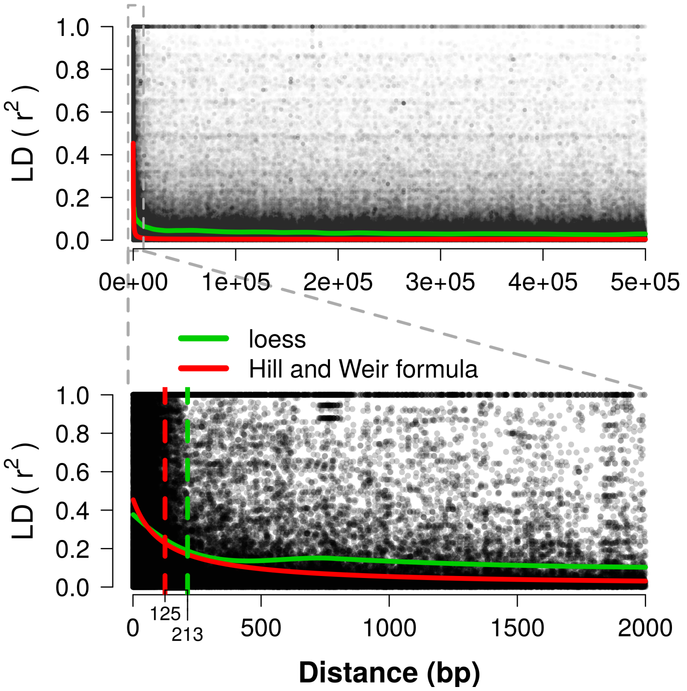


**Figure S10** Linkage disequilibrium decay of the B1K+ collection across genome zooming in the first 500 kb and 2 kb. The green and red lines represent the fitted $r^2$ values by using *loess* method and Hill and Weir formula. The green and red dash lines represent the decay to the half of the highest fitted $r^2$.

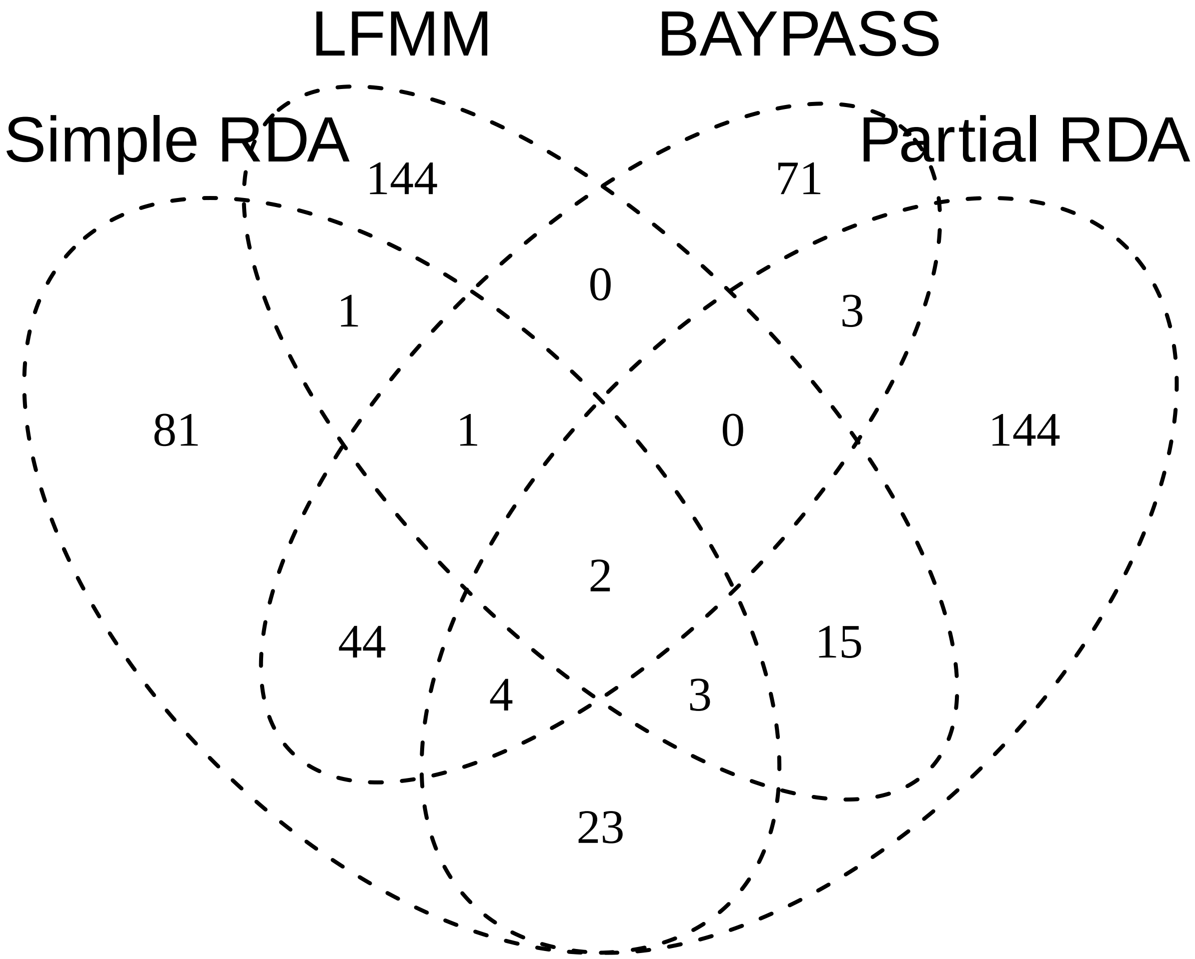


**Figure S11** Number of genes locating within 500 bp adjacent intervals of significant SNPs detected by the GEA and outlier methods.

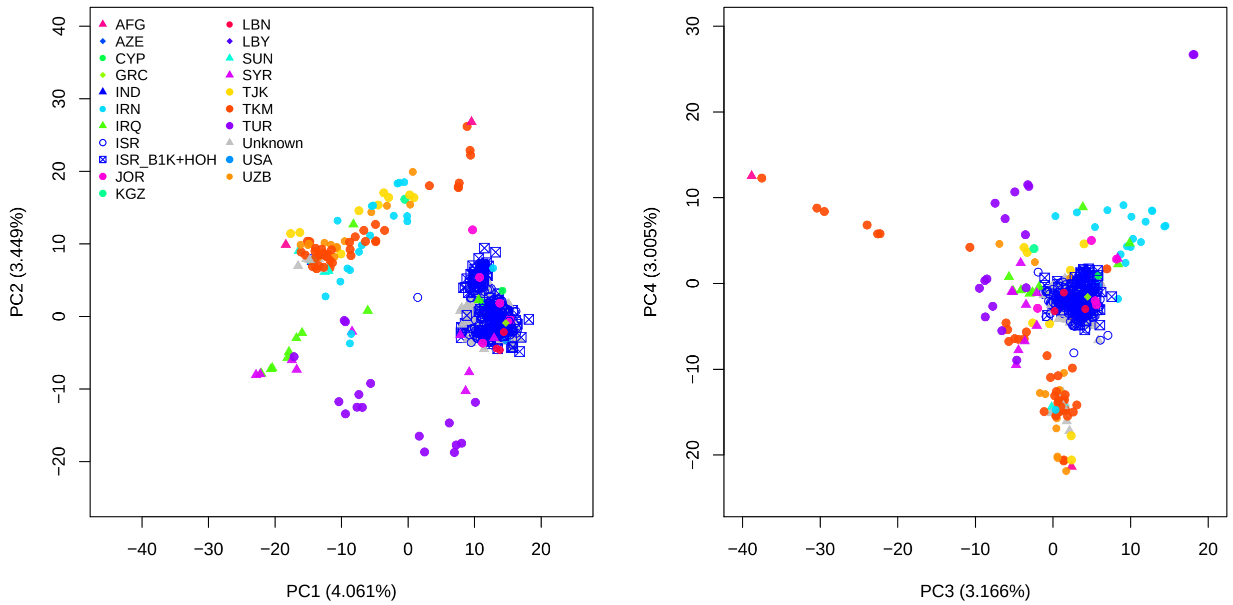


**Figure S12** PCA plots of 244 B1K+ accessions and 1,121 accessions from IPK's genebank with countries of origins. (A) Result of PCA performed by using all of the available accessions. (B) Result of PCA performed by using 72 geographically diverse accessions and projecting the remaining accessions to PC spaces. The 244 B1K+ accessions are represented by blue boxes with crosses inside. The blue open dots represent IPK accessions originating in Israel and the gray closed triangles represent IPK accessions with unknown origins. The country abbreviations are according to ISO 3166-1 $\alpha$-3.

#### Supplementary Files

**File S1** Geographical coordinates and environmental data of 244 B1K+ used in this study. Both raw environmental data and twelve environmental variables used in genome-environment association analysis are included in the file. Grouping information of 58 demes and 10 manually defined regions used in gene flow analysis are also included.

**File S2** A demonstration showing the influence of marker number and additional samples (HOH accessions) on the result of population structure analyses.

**File S3** Statistics of the genome-wide scan methods for 27,147 SNPs, including $X^T X$ of *BAYPASS*, and the $p$ and $q$ values of the simple RDA, partial RDA and LFMM.

**File S4** Annotations of genes in the upstream or downstream 500 bp of the candidate SNPs detected by the genome scan methods (*BAYPASS*, simple RDA, partial RDA and LFMM).
